# Long term analysis of social structure: evidence of age-based consistent associations in male Alpine ibex

**DOI:** 10.1101/2021.12.02.470954

**Authors:** Alice Brambilla, Achaz von Hardenberg, Claudia Canedoli, Francesca Brivio, Cédric Sueur, Christina R Stanley

## Abstract

Despite its recognized importance for understanding the evolution of animal sociality as well as for conservation, long term analysis of social networks of animal populations is still relatively uncommon. We investigated social network dynamics in males of a gregarious mountain ungulate (Alpine ibex, *Capra ibex*) over ten years focusing on groups, sub-groups and individuals, exploring the dynamics of sociality over different scales. Despite the social structure changing between seasons, the Alpine ibex population was highly cohesive: fission-fusion dynamics lead almost every male in the population to associate with each other male at least once. Nevertheless, we found that male Alpine ibex showed preferential associations that were maintained across seasons and years. Age seemed to be the most important factor driving preferential associations while other characteristics, such as social status, appeared less crucial. We also found that centrality measures were influenced by age and were also related to individual physical condition. The multi-scale and long-term frame of our study helped us show that ecological constrains, such as resource availability, may play a role in shaping associations in a gregarious species, but they cannot solely explain sociality and preferential association that are likely also to be driven by life-history linked physiological and social needs. Our results highlight the importance of long-term studies based on individually recognizable subjects to help us build on our understanding of the evolution of animal sociality.

## Introduction

Understanding the drivers of spatial and temporal interactions between animals is of great importance to some of the most pressing questions in biology, such as understanding the evolution of animal sociality (Couzin & Laidre, 2009), determining how genetic and cultural information spread within and among inter- connected populations (Van de Waal & Bshary, 2011), tracking disease transmission (MacIntosh et al., 2012; Marchand et al., 2017; Silk et al., 2019), and predicting the invasion dynamics of introduced species (Fogarty et al., 2011).

In many social species, the process of group formation is highly dynamic, with frequent changes in group size and composition (Grueter et al., 2020). Groups can merge (fusion) or split (fission) over time and space, making group composition a dynamic property (Sueur et al., 2011). Social systems characterized by fission-fusion dynamics are widespread among vertebrates as for example fish (*Poecilia reticulata*, Wilson et al., 2014), birds (Silk et al., 2014; Papageorgiou & Farine, 2020) and several mammals including primates (Smuts et al., 1987), Bechstein’s bats (*Myotis bechsteinii*, Kerth & König, 1999), bottlenose dolphins (*Tursiops spp*., Connor & Wells, 2000), spotted hyenas (*Crocuta crocuta*, Holekamp et al., 1997), African elephants (*Loxodonta africana*, Wittemyer et al., 2005), giraffes (*Giraffa camelopardalis*, Bond et al., 2019), feral goats (*Capra hircus*, Stanley & Dunbar, 2013) and sheep (*Ovis aries*, Jewell et al., 1974). The timescale over which fission-fusion dynamics occur may differ as environmental conditions or individual requirements change (Aureli et al., 2008; Sueur et al., 2011). Indeed, there is pronounced variation in the degree of fission-fusion dynamics both across and within species which can vary from a highly cohesive society with stable group membership to a highly fluid society with either relatively stable or flexible subgroup membership (Aureli et al., 2008). The flexibility of fission-fusion dynamics is likely to have evolved because it allows individuals to optimize the costs and benefits of group-living (Moscovice et al., 2020). Fission-fusion dynamics are predicted to be most frequent in environments with spatial variability and to increase with temporal uncertainty and unpredictability of the environment (Sueur et al., 2011). Other factors known to influence the structure and cohesion of groups are demographic processes such as deaths, births, dispersal and immigration (Ilany & Akçay, 2016) that may influence social structure through the loss of some social connections and the formation of new ones (Shizuka & Johnson, 2020) as well as by driving changes in patterns of association between remaining individuals (e.g., Flack et al., 2006).

Studies on social systems often focus on the association between individuals, as members of a group can show a preference to associate with specific individuals and maintain certain social links despite the frequent splitting and merging of the overall group. In many social mammals there is a tendency towards homophily, i.e., for individuals to associate with others that share similar characteristics such as age or sex (Le Pendu et al., 1995). A possible explanation for these preferences is that the cost of living in groups with members of the same age class or sex is lower due to similar physiological needs and a reduction in harmful inter or intra-sexual interactions (Conradt & Roper, 2000; Ruckstuhl & Neuhaus, 2000). Likewise, social relationships between individuals may be affected by kinship (Cassinello & Calabuig, 2008; Wittenmyer et al., 2009; Podgórski et al., 2014) as well as by social rank as observed in horses (*Equus caballus*, Kimura, 1998) and red deer (*Cervus elaphus*, Appleby, 1983), where individuals spent more time close to other of similar rank.

Associations can be stable or can change during the life on an individual, varying from long-term or even life-long associations (as shown, e.g., in birds, Teitelbaum et al., 2017; wild boars, *Sus scrofa*, Podgórski et al., 2014; feral horses and ponies, Cameron et al., 2009 and Stanley et al., 2018) to transitory or short-term associations (e.g., in spotted hyenas, Smith et al., 2011). Also the levels of gregariousness can differ between individuals of the same species or population and can be influenced by various factors such as age (Carter et al., 2013; Machanda et al., 2020) or reproductive status (Fischhoff et al., 2009; Vander Wal et al., 2015; Machanda & Rosati, 2020). This can, however, also be a consistent individual characteristic that is maintained throughout life as recent studies on personality (i.e., the presence of behavioural differences that are repeatable over time and across situations) have shown (e.g., Krause et al., 2010; Firth et al., 2018). Sociability is indeed one of the five personality traits described in literature and it can affect reproductive success and survival and hence fitness (Réale et al., 2007).

Social network analysis (SNA) is a powerful tool that can be applied to the analysis of several aspects of social behaviour (Sueur et al 2011; Sosa et al., 2021b). One of the most attractive features of SNA is that it allows to study the social organisation of animals at all levels (individual, dyad, group, population) and for all types of interaction (e.g., aggressive, cooperative, sexual, Krause et al., 2009), allowing a plethora of novel insights into the evolution and maintenance of sociality to be elucidated (Wey et al., 2008; Pinter-Wollman et al., 2014; Krause et al., 2015; Webber and Vander Wal 2019; Sosa et al., 2021b). As social structure can also affect population growth rates, dispersal and gene flow, network analysis also has the potential to be an important tool in the management of wild populations (Tarlow & Blumstein, 2007; Schakner et al., 2017, Snijders et al., 2017, Welch et al., 2020).

To date, few studies have investigated dynamic changes in social network structures of wild populations in the long term, especially where the individuals composing the network change due to demographic processes (e.g., death, emigration or immigration of individuals, Cantor et al., 2012; Borgeaud et al., 2017). Shizuka & Johnson (2020) called for the need to integrate demographic processes and consequent social processes into social network analyses. However, the scarcity of such studies is probably due to the intensity of consistent data collection required in order to incorporate environmental and demographic changes or stochastic events, in addition to limitations in the available analytical tools (Pinter-Wollman et al., 2014). Instead, many studies making use of SNA in free ranging populations rely on relatively short-term datasets and, therefore, are only able to describe snapshots of the social systems. Since long term monitoring of wild species has provided important contributions to the study of ecology and evolution as well as to conservation (Festa-Bianchet et al., 2017, Clutton-Brock, 2021), long term analysis of social network dynamics could bring rich rewards in terms of a better understanding of population-level processes (Pinter-Wollman et al., 2014).

Ungulate species are widespread in all continents except Antarctica and show high diversity both in terms of size, habitat as well as behaviour (Wilson & Mittermeier, 2011). Their social organization has been extensively studied and provided crucial elements for understanding the evolution of vertebrates’ social systems (Jarman, 1974, Krause et al., 2002). However, many of the studies on social networks of ungulates have focused on female associations (Vander Wal et al., 2016; Ramos et al., 2019) meaning knowledge on male sociality in ungulates remains scarce. To fill those gaps, we performed a long-term analysis of male social structure in a gregarious ungulate: the Alpine ibex (*Capra ibex*). We took advantage of a detailed long-term dataset resulting from ten years of behavioural observations of individually identifiable male Alpine ibex to explore the male social network of a gregarious ungulate and to investigate population-level changes in social structure over time.

The Alpine ibex, a mountain ungulate of conservation concern (Brambilla et al., 2015; 2018; 2020), is a gregarious species that lives in open-membership groups and shows fission-fusion dynamics. Alpine ibex are highly sexually dimorphic and exhibit strong sexual segregation all year round except for during the mating season, that occurs between December and January (Villaret & Bon, 1995; Ruckstuhl & Neuhaus, 2001). As the degree of sexual segregation increases with male age (Ruckstuhl & Neuhaus, 2001), outside the rut, adult males join groups mainly composed of males, while yearlings stay in groups mainly composed of females, and young males (≤ 2-3 years old) may move between male and female groups.

This study was conducted during spring and summer months, when sexual segregation is at its peak, and focused on the social network of male Alpine ibex. Specifically, we aimed to determine: a) whether males showed consistent associations and which factors influenced the choice of preferred companions; b) which individual characteristics (age, life stage) influenced network metrics at the individual level and if these metrics were consistent over time; c) whether social structure was stable across seasons (i.e., across periods with different ecological conditions); d) whether any changes in the social network structure over time could be explained by demographic factors.

Similarly to observations in other bovids (Vander Wal et al., 2015; Ramos et al., 2019), we expected to find a highly connected and cohesive social network, with all the individuals connected either directly or indirectly. Despite the aforementioned studies having analysed social structure in females, based on field observations we expected similar results also for males. Age was expected to predict certain network node-based measures (Turner et al., 2018; Sosa et al., 2021b). Particularly, we expected adult males to occupy the more central positions in the network due to their reproductive status and generally higher rank. In addition, we expected males to become less central once they had passed their reproductive peak due to their decreased competitive ability. Age often underpins social preferences (e.g., Wey & Blumstein, 2010; Welch et al., 2020); we therefore expected stronger associations between males of the same age class due to their similarity in body size, social motivation, and behaviours. At the same time, and as rank-based homophily was observed in other species (Sosa, 2016), we also expected stronger associations between males with similar social rank.

At the population level, we wanted to verify if the structure of the social network remained stable between seasons and between years over a period of ten years. Environmental conditions during the study period were reasonably stable within seasons so we did not expect variations in global network measures. However, as more experienced, older individuals are expected to help maintain group cohesion (Allen et al., 2020), we expected to find a correlation between global centrality measures and the proportion of old males in the population which could change due to stochastic events (e.g., harsh winters).

## Methods

### Study area and population

The study was conducted on a free-ranging population of Alpine ibex in the Levionaz basin (Valsavarenche valley, AO), within the Gran Paradiso National Park (North-western Italian Alps - 45°26′ N, 7°08′ E).

In the study site, ibex are captured and individually marked with coloured plastic ear tags (Allflex®: Allflex Europe (UK), 77 Greenchurch Street, London) in the framework of a long-term study on the life-history and conservation of the species (Bassano et al., 2003; Bergeron et al., 2010; Brambilla & Brivio, 2018). The capture and marking protocol used in this study has been authorised by the Italian Ministry of Environment (authorisation nr. 25114 of 21/09/2004) after the positive review by the Italian National Institute for Environmental Protection and Research (ISPRA) and was developed trying to minimize the effects on the welfare of the animals (Brambilla et al., 2013; Brivio et al., 2015).

The mean number ± SD of individuals counted in the Levionaz population during the 10 years of this study was 180 ± 41 of which 63.9 ± 10.8 were males (table 6). Censuses were conducted every year in July block-counts by the personnel of Gran Paradiso National Park. The proportion of marked males varied from a minimum of 50% in 2017 to a maximum of 88% in 2011 and 2012 for a total of 111 marked individuals observed during the study.

### Data collection

Data were collected for male Alpine ibex during a period of ten years from 2008 to 2017 in spring (May-June) and summer (July-Aug-Sept). As Alpine ibex perform yearly altitudinal migrations following the green up of the vegetation (Parrini et al., 2003), exact dates of the seasonal ranges varied between years and were selected according to the altitudinal movement of the animals on the study site.

#### Individual attributes

*Age*. The age of the marked individuals was determined at capture by counting the number of annual horn segments (Brambilla & Canedoli, 2014). The age class of the unmarked individuals was estimated by using binoculars to count the number of annual horn segments. Unmarked individuals were divided in age classes, defined as follows based on clearly visible body mass differences (Couturier, 1962): 2-5 years old (young); 6-8 years old (sub-adults); 9-11 years old (fully grown adults); >11 years old (old individuals). *Season preceding death*. Male Alpine ibex adult survival is very high and most of them reach senescence (ToÏgo et al., 2007). However, some individuals die earlier and, particularly in case of chronic diseases, their behaviour could change in the months preceding death. We therefore recorded all deaths of identified individuals to obtain a binary variable “season preceding death”; this indicated whether the animal died within that year of data collection or if it survived until the following year. This was possible as carcasses of dead individuals were often found in the field within a few days of death. When possible, the cause of death was also ascertained. Some individuals’ carcasses, however, were not found. This happened especially when animals died during winter or in an inaccessible part of the study area. If an animal was not observed in the study site nor in the surrounding areas (monitored daily by park rangers) for a whole year, we considered it as dead. We could not exclude that some of those individuals migrated; however, as all the animals included in our study were observed several times every year and their movements were rather predictable, we were confident in considering them as dead if they were not observed again. Indeed, during the time of our study, we had no cases of animals that were observed again in our or in other study areas after one whole year when they were not observed.

#### Male associations

Association patterns between individuals were defined via the ‘gambit of the group’ method (Whitehead, 2008), i.e., assuming that each animal in a group is associating with every other individual in that group (Croft et al., 2008; Franks et al., 2010). Association data were collected daily by means of surveys conducted when the animals were active/feeding. When possible, associations were recorded twice per day, after dawn and before dusk, considering each survey as independent due to the strongly bimodal daily activity pattern typical of the species, but time and number of daily observations varied based on seasonal and weather conditions (details on the number of surveys per season are provided in supplementary material S0). Data were collected by an observer scanning the study area while walking along transects consisting of the GPNP paths used during block counts. Transect routes changed during the season according to the altitudinal movement of the animals in the study site (Parrini et al., 2003). Each session of data collection lasted around one to three hours depending on the location of the animals. Every year, data were collected by two to four observers. New observers were trained and tested by the same person that was present for the whole duration of the study. Before new observers could start collecting data, blind contemporary data collection with the expert observer was conducted for up to one week, until consensus in group identification was reached. For each group, the total number of individuals, identity of the marked individuals and age class of the unmarked individuals were recorded. Two groups were considered as distinct if their closest members were more than 50 metres apart. This threshold was set after field observations considered this distance sufficient to avoid social interactions during random movements while foraging. Any animal observed alone was considered to be a separate group. Data on association were used to build association matrices for each season of each year (see the data analysis paragraph for details).

#### Male dominance

All observed agonistic interactions when an individual was clearly dominant over another, as defined by Bergeron et al., (2010), were recorded using all occurrence sampling (Altmann, 1974). Agonistic interactions were collected opportunistically during the group composition data collection as well as during focal samples conducted for other studies. The identity of the winner and loser in each interaction was recorded, allowing dominance matrices to be built based on the frequency of dominance interactions per dyad. If the identity of one or both of the individuals involved in the interaction was unknown, this interaction was not included in the matrix.

The outcome of agonistic interactions was also used to calculate hierarchical rank via the Elo-rating method (Elo, 1978; Albers & De Vries, 2001, Neumann et al., 2011). This method is based on the sequence in which interactions occur and continuously updates ratings by looking at interactions sequentially (Neumann et al., 2011). As previous analysis showed that hierarchical ranks are established early in spring and remain rather stable during summer, we calculated the Elo score (R package EloRating, Neumann et al., 2011) for each individual as of the 30^th^ of June of each year in which the individual was observed.

### Network analysis

#### Association networks

The association network was built based on the half weight index (HWI; Whitehead, 2008). HWI is defined as:

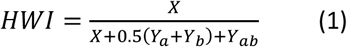

where *X* is the number of sampling periods during which individuals a and b were observed together, *Y*_*a*_ is the number of sampling periods when *a* was observed without *b, Y*_*b*_ is the number of sampling periods in which b was observed without a, and *Y*_*ab*_ is the number of sampling periods when *a* and *b* were both observed separately (Whitehead, 2008). The HWI is considered to be less biased when not all associating individuals can be identified (Whitehead, 2008) and it was chosen as association index to account for the unequal detection probability typical of gregarious mountain ungulates (Vander Wal et al., 2015) and for potential uneven sampling between individuals. As ecological conditions (e.g., resource quality and distribution, temperatures, precipitations) change during the year and Alpine ibex perform seasonal altitudinal migrations (Parrini et al., 2003), the HWI for each pair of individuals was calculated separately for spring and summer of each year (2008-2017) resulting in 20 time-aggregated association matrices (for details on the methods and tools used to build the networks, see the data analysis section).

#### Dominance networks

Dominance networks were built using the absolute frequency of dominance interactions between each pair of individuals within each season and year (as we assume that our method allowed random sampling of dominance interactions), resulting in 20 dominance networks. Dominance networks were non-symmetrical as the directionality of the dominance resulting from the interaction was preserved.

#### Network metrics

To analyse the network structure of male Alpine ibex, we used the metrics described below.

*Individual-level metrics: strength centrality* and *eigenvector centrality* were calculated for each individual in each network and averaged among individuals for each network. *Strength centrality* is calculated as the sum of the weights of the edges in a weighted network (Wasserman & Faust, 1994). This measure represents the sociality of an individual as it estimates the frequency of its interactions (Sosa et al., 2021a) and it was also used to compute gregariousness at the group level (see *Global metrics* section). *Eigenvector centrality* is defined as the first non-negative eigenvector value obtained by transforming an adjacency matrix linearly (Wasserman & Faust, 1994). It measures the centrality of a node by examining its connections as well as that of its alters (with alters being the other individuals connected to the node). Eigenvector centrality can therefore be interpreted as the social resources available to an individual (Brent et al., 2011).

*Global metrics*: as measures to describe the global structure of the network we calculated *network density, gregariousness* and *Typical Group Size (TGS). Network density* calculates the ratio between existing links and all potential links of a network, and it assesses the connection of the network as a whole (Sosa et al., 2021a). *Gregariousness* represents the tendency of individuals to associate with few or many individuals (Godde et al., 2013). At the individual level, it is represented by *strength centrality* and it is calculated as the sum of the values of the association indices involving that individual. The overall gregariousness of a population is calculated as the average gregariousness of all individuals in the population. *TGS* quantifies group size as experienced by an average individual of the population and it emphasises the extent to which members of the population tend to associate (Jarman, 1974). It is an animal-centred measure defined as the sum of the squares of the number of individuals in each group, divided by the total number of animals sampled (Jarman, 1974).

### Data analysis

Association matrices were built in SOCPROG 2.9 (Whitehead, 2009). Network analysis was performed in R (R Core Team, 2020) using the packages tnet (Opsahl, 2009), network and sna (Butts, 2020), included in the statnet suite (Statnet Development Team, 2003-2020; Handcock et al., 2008) and with the package ANTs (Sosa et al., 2020). Generalised Linear Mixed Models (GLMMs) were conducted using the R packages ANTs and lme4 (Bates et al., 2015). Details of which tool was used for each analysis are provided below.

Data and code are accessible on the Dryad Digital Repository with the following DOI https://doi.org/10.5061/dryad.w0vt4b8st

#### Association patterns

To investigate association patterns, we used a Quadratic Assignment Procedure (QAP) approach to compare pairs of association networks. The QAP test is a specialised version of the Mantel test that uses random permutations of node labelling to determine whether a correlation between two matrices is significantly higher than expected (Krackhardt, 1988; Krause et al., 2015). QAP tests were carried out with 10 000 permutations using the sna package.

To test for consistency in associations between seasons, a QAP test was carried out between the two seasonal association networks within each year. For this analysis, the networks were built excluding individuals that were not observed across both seasons within a year.

Furthermore, to test for consistency in associations between years, we compared the networks built for the summer seasons of the years 2012-2016. The network used for this analysis only included a subset of individuals that were observed in all those five years (hereafter referred to as subset networks). This time period was selected as being optimal in terms of having the largest number of individuals present across the entire period, hence allowing the greatest power for testing for consistency in associations; the summer seasonal network was selected as spring associations may partly be driven by resource availability. Moreover, spring and summer networks within each year were correlated (see Results section). QAP tests were carried out between each subset network and the subset network of the following year.

To test whether the age difference between pairs of individuals had an impact on whether they were likely to associate or to be dominant over each other, we built matrices of difference in age (in years) within each dyad for each year. We built both a symmetrical matrix with absolute age difference as well as an asymmetrical matrix with the exact age difference between dyads. We then used QAP tests to determine whether there was significant consistency in structure between the summer association matrix and the absolute age difference matrix, then between the dominance matrix and the age difference matrix. Finally, to test whether dominance rank differences between individuals predicted their likelihood of association, we built matrices of absolute difference in their Elo scores (used as a proxy for hierarchical rank) and performed a QAP test to test for structural consistency between the summer association matrix and the absolute Elo score difference matrix. As this resulted in multiple hypothesis testing, for each of the sets of QAP tests described before, we applied a sequential Bonferroni correction (Holm, 1979) for the assessment of significance levels. A combined *p-*value for all years was calculated using Fisher’s method with the package poolr (Cinar & Viechtbauer, 2020).

#### Factors affecting node measures and seasonal network structure

To determine whether association network structure changed between seasons (i.e., between periods with different ecological conditions) and which individual characteristics predicted node-based measures, we carried out Generalized Linear Mixed Models (GLMMs) on permuted association matrices with the package ANTs: time-aggregated networks for each season of each year were built through data stream permutations resulting in a list of 20 symmetrical association matrices (each with 10 000 permutations) used to calculate the centrality measures of interest (strength and eigenvector centrality) and to run permuted GLMMs.

The fixed and random structures of the permuted GLMMs were selected using non-permuted GLMMs (lme4 package): each GLMM included as the dependent variable either strength centrality (modelled with a gaussian distribution) or eigenvector centrality (modelled with a binomial distribution) and, as fixed effects, the season and individual characteristics: age as a quadratic term (Bergeron et al., 2008) and the season preceding death (as a binary variable that indicated whether the animal died within that year of data collection or if it survived until the following year). The models also included year and individual identity (ID) as random effects. The choice of the fixed effects as well as the assessment of the importance of individual ID as a random effect was made based on the Akaike Information Criterion (AIC, Akaike, 1973; Burnham et al., 2011), with Δ AIC>2 used as a threshold for the selection of the best fitting models. Graphical analysis of the residuals and the coefficient of determination R^2^ were used to check model fit. Both marginal (R^2^m, that describes the proportion of variance explained by the fixed factors alone) and conditional coefficient of determination (R^2^c, which describes the proportion of variance explained by both the fixed and random factors) were calculated with the R package MuMIn (Barton, 2009).

#### Seasonal and annual changes in the social structure

To investigate seasonal differences in the social structure, we run linear models to compare the spring and summer values of the global measures (mean gregariousness and TGS, calculated in R with own-built functions). To account for the sampling effort, the number of surveys for each season was added as an explanatory variable to the model.

To detect possible changes in the network structure during the ten years of the study, we calculated the global measures of density (calculated within the network R package), gregariousness and TGS within each year (across seasons). As we hypothesised that the age structure of the population could affect the cohesion of the social structure and hence the above-mentioned global network measures, we calculated the proportion of adult individuals (i.e., of individuals of 9 years and more) and the total number of males in the population (counting the maximum total number of males, including both marked and unmarked individuals, observed during the daily observations). We then ran two separate generalised linear models (GLM) with binomial distribution to test the effect of the proportion of old individuals and of the total number of males on density. In addition, we ran three GLMs to test the effect of the proportion of old individuals and of the total number of males on TGS and to test the effect of the total number of males on gregariousness.

## Results

The association networks of Alpine ibex showed a high level of connectivity (e.g., figure 1) with the network density being close to one in all the years of the study.

**Figure 1.**
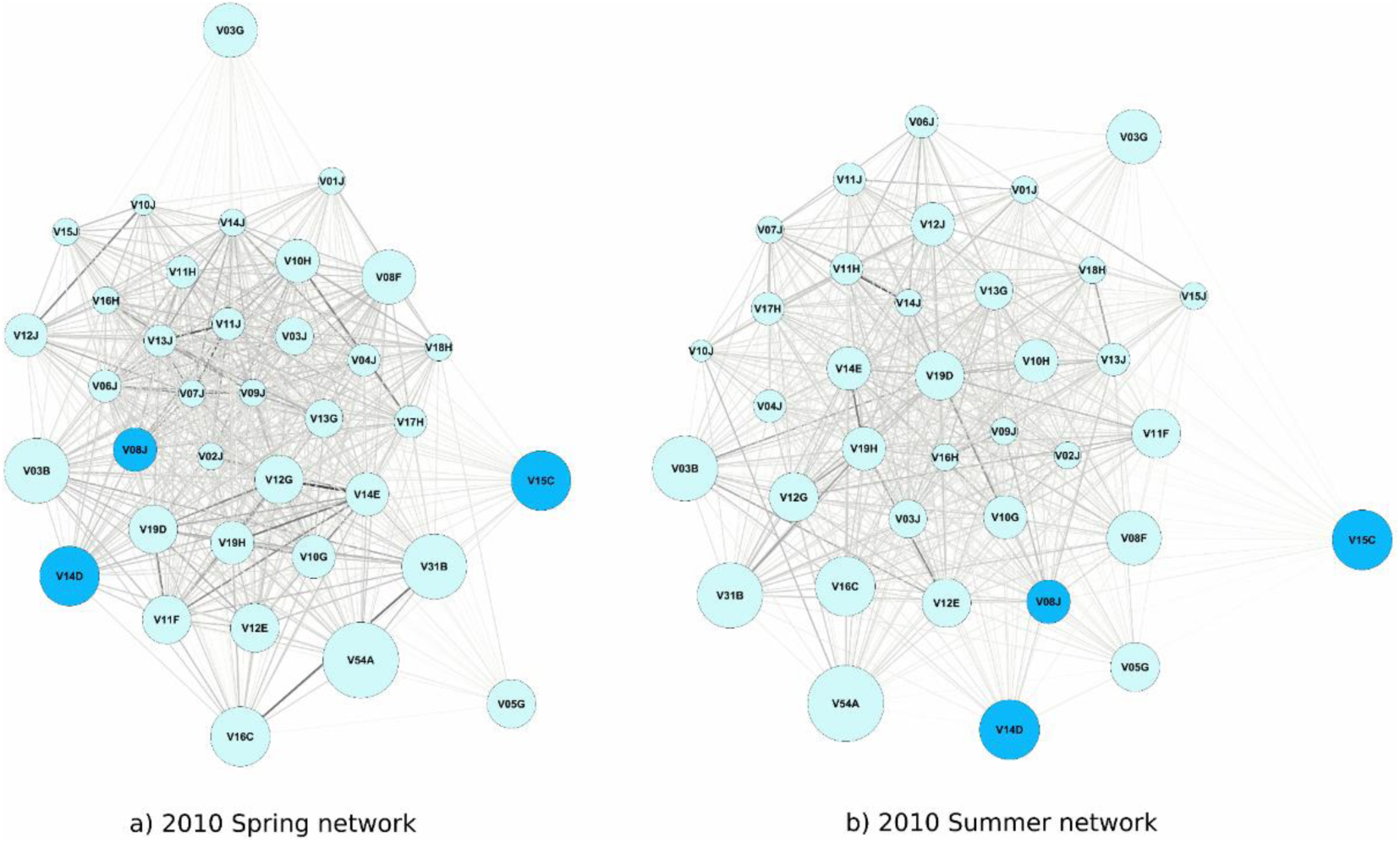
An example of a weighted association network of male Alpine ibex in a) spring and b) summer of the same year (2010). Graphs were created using Gephi 0.9.2 (Bastian et al., 2009) with the “ForceAtlas” spatialization tool. The colours and width of the edges are proportional to the strength of association (with darker and thicker edges representing stronger associations). The size of the nodes represents the age of the individuals. The colours of the nodes represent the value of the binary variable “season preceding death” of the individuals during the year. Individuals that die in the months following data collection are represented in darker colour. The graphical visualization of all networks (all season of all years) can be found in Supplementary Materials S1.

### Association patterns

QAP tests performed to test for consistent associations across seasons within each year showed significant correlations between association matrices within all the ten years of the study (table 1) indicating that male Alpine ibex maintain consistent bonds throughout the year, despite changes in environment and location. Consistent associations between adjacent years were also observed over a five year period as shown by the results of the pairwise QAP tests performed on the subset networks between summer association networks in adjacent years (table 2).

**Table 1.** Results of QAP tests to investigate consistency in association network structure across the spring and summer seasons within each year. Significant correlations after a Bonferroni sequential correction are indicated by bold *p-*values. Combined *p*-value < 0.001.

| Seasons | Correlation coefficient | *p*-value |
| --- | --- | --- |
| Spring vs summer 2008 | 0.250 | **0.004** |
| Spring vs summer 2009 | 0.511 | **<0.001** |
| Spring vs summer 2010 | 0.526 | **<0.001** |
| Spring vs summer 2011 | 0.490 | **<0.001** |
| Spring vs summer 2012 | 0.407 | **<0.001** |
| Spring vs summer 2013 | 0.315 | **<0.001** |
| Spring vs summer 2014 | 0.236 | **0.016** |
| Spring vs summer 2015 | 0.460 | **<0.001** |
| Spring vs summer 2016 | 0.452 | **0.007** |
| Spring vs summer 2017 | 0.493 | **<0.001** |

**Table 2.** Results of QAP tests to investigate consistency in association network structure between summer seasons of adjacent years. Tests were performed on subset networks including N=21 individuals that were observed in all the years 2012-2016. Significant correlations after a Bonferroni sequential correction are indicated by bold *p-*values. Combined *p*-value < 0.001.

| Years | Correlation coefficient | *p*-value |
| --- | --- | --- |
| 2012 vs 2013 | 0.236 | **0.014** |
| 2013 vs 2014 | 0.192 | **0.028** |
| 2014 vs 2015 | 0.265 | **0.016** |
| 2015 vs 2016 | 0.400 | **0.008** |

We also found that dominance matrices mostly correlated positively with age difference matrices within the summer season for each year (table 3a), indicating that dominance interactions were more likely to occur between individuals the greater their difference in age, with the older individual of the dyad being more likely to be dominant over the younger. Association matrices, instead, mostly correlated negatively with their associated absolute age difference matrices, showing that individuals were more likely to associate with those closest in age (table 3b). Finally, the tests between association matrices and absolute difference in Elo score matrices showed no evidence of a correlation between association choice and dominance rank differences or similarities (table 3c).

**Table 3.**
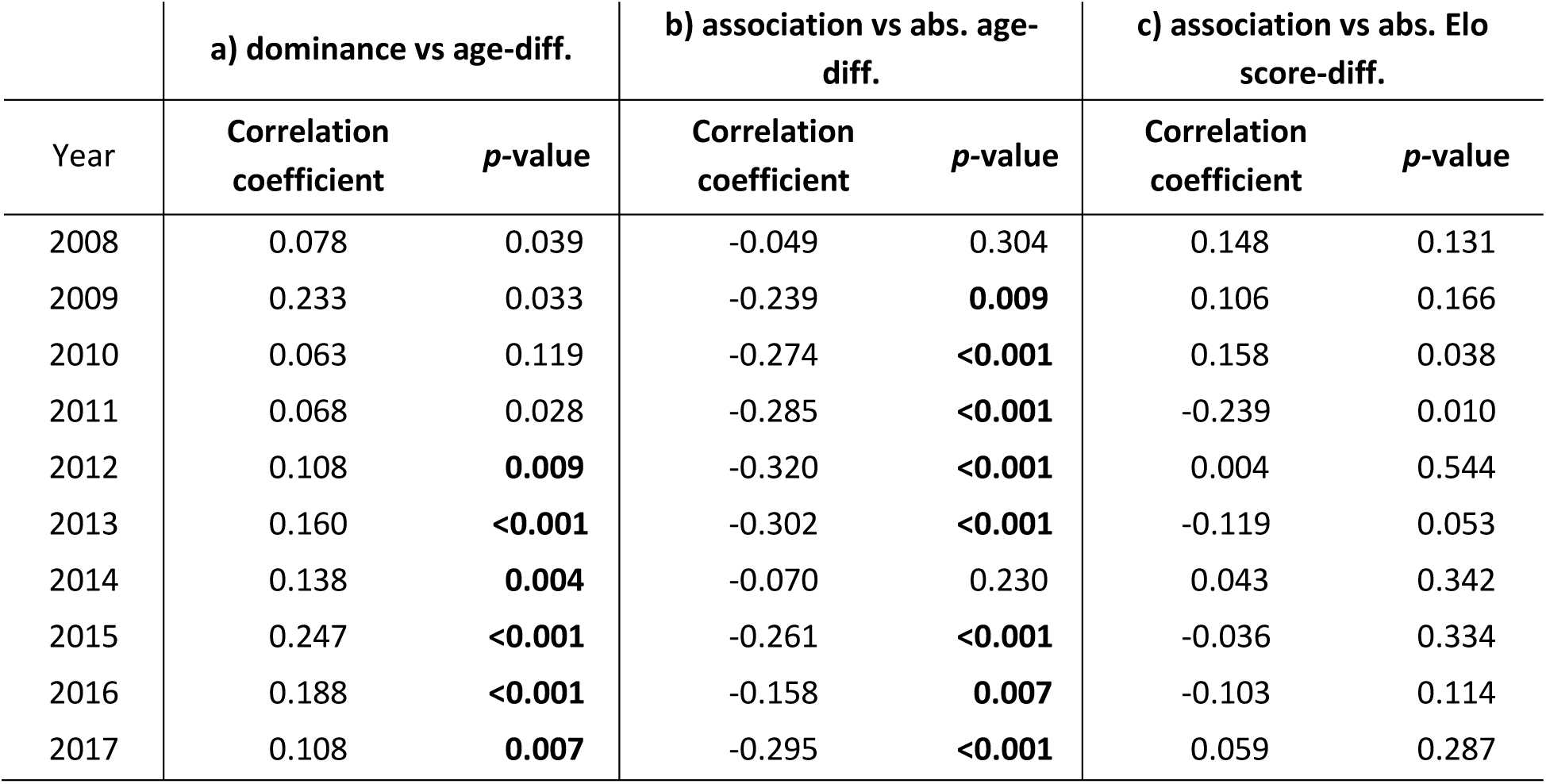
Results of QAP tests for correlations between a) dominance matrices and their associated exact age difference matrices (combined *p-*value <0.001); b) association matrices and their associated and absolute age difference matrices (combined *p-*value <0.001); c) association matrices and their associated absolute Elo score difference matrices. QAP tests were performed only on the summer season matrices. Significant correlations after a Bonferroni sequential correction are indicated by bold *P* values.

### Factors affecting node measures and seasonal network structure

The results of the GLMMs, performed to test whether the association network structure changed between seasons and which individual characteristics predicted node-level centrality measures, are presented in tables 4 and 5. Model selection results are presented in the supplementary materials S2.

**Table 4.**
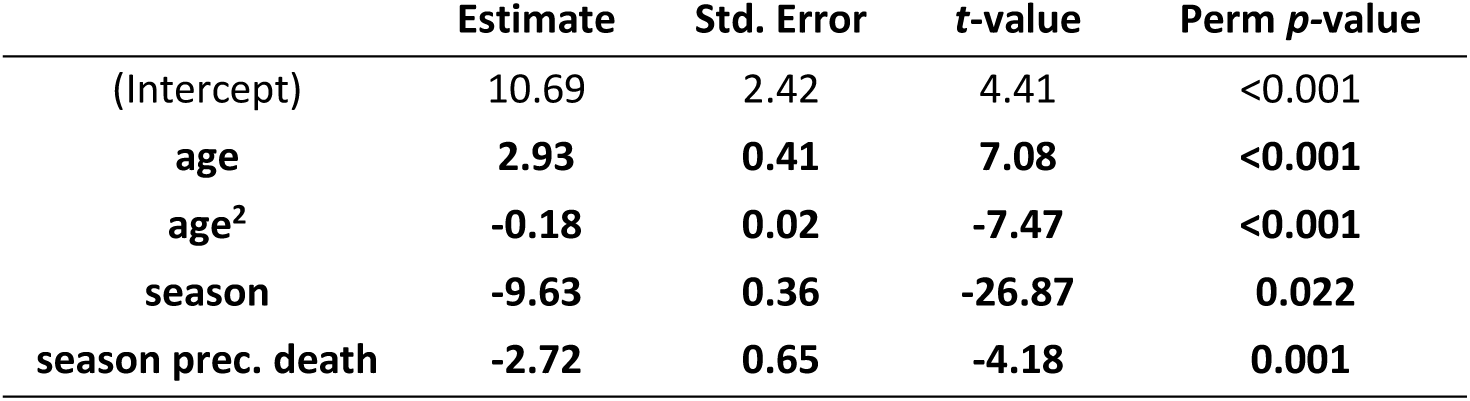
Output of the permuted GLMM performed to explain the variance of strength centrality. Perm *p-* value represents the permuted *p-*values obtained after 10 000 permutations in ANTs. Model specification: strength ∼ age + age^2^ + season + season preceding death + (1|year) + (1|ID), family=gaussian.

| | Estimate | Std. Error | t-value | Perm $p$-value |
| --- | --- | --- | --- | --- |
| (Intercept) | 10.69 | 2.42 | 4.41 | <0.001 |
| age | **2.93** | **0.41** | **7.08** | **<0.001** |
| age<sup>2</sup> | **-0.18** | **0.02** | **-7.47** | **<0.001** |
| season | **-9.63** | **0.36** | **-26.87** | **0.022** |
| season prec. death | **-2.72** | **0.65** | **-4.18** | **0.001** |

**Table 5.**
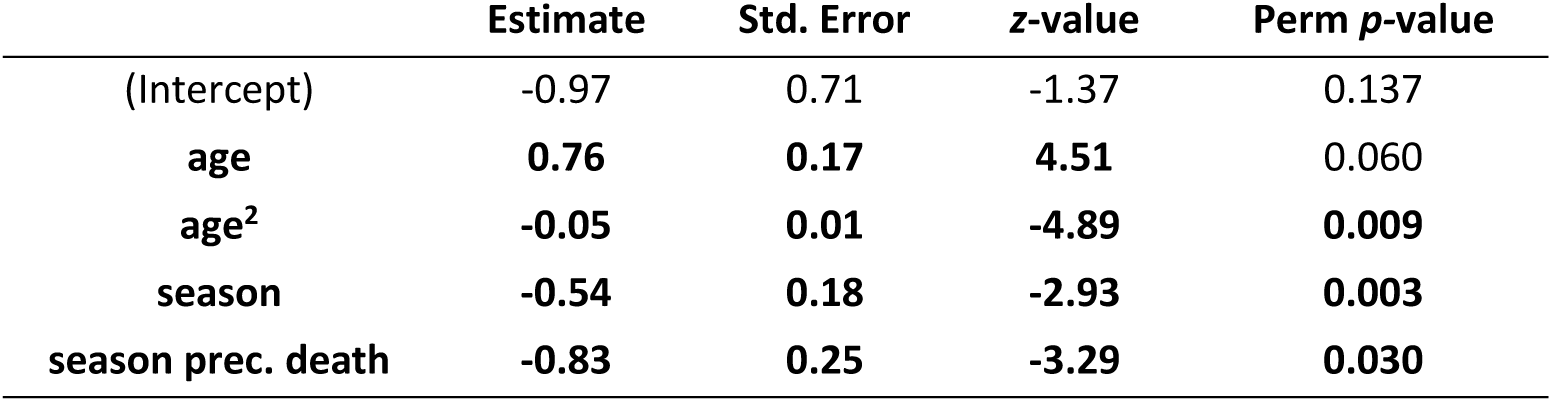
Output of the permuted GLMM performed to explain the variance of eigenvector centrality. Perm *p-*value represents the permuted *p* values obtained after 10 000 permutations in ANTs. Model specification: eigenvector ∼ age + age^2^ + season + season preceding death + (1|year) + (1|ID), family= binomial.

Strength centrality was significantly lower in the summer season compared to spring and was correlated with age following a quadratic curve, i.e., it increased with age, until around 9-10 years, but then decreased when animals became older (table 4, figure 2). Strength centrality was also negatively correlated with the season preceding death (i.e., the occurrence of death in the months following the observations). In the best fitting model, ID was retained as a random factor (the ΔAIC between the selected model with and without ID as random factor was 134.1) and accounted for almost 14% of the variance explained by the model (13.88 ± 3.72). The coefficients of determination of the selected model were: R^2^m=0.30, R^2^c=0.72.

**Figure 2.**
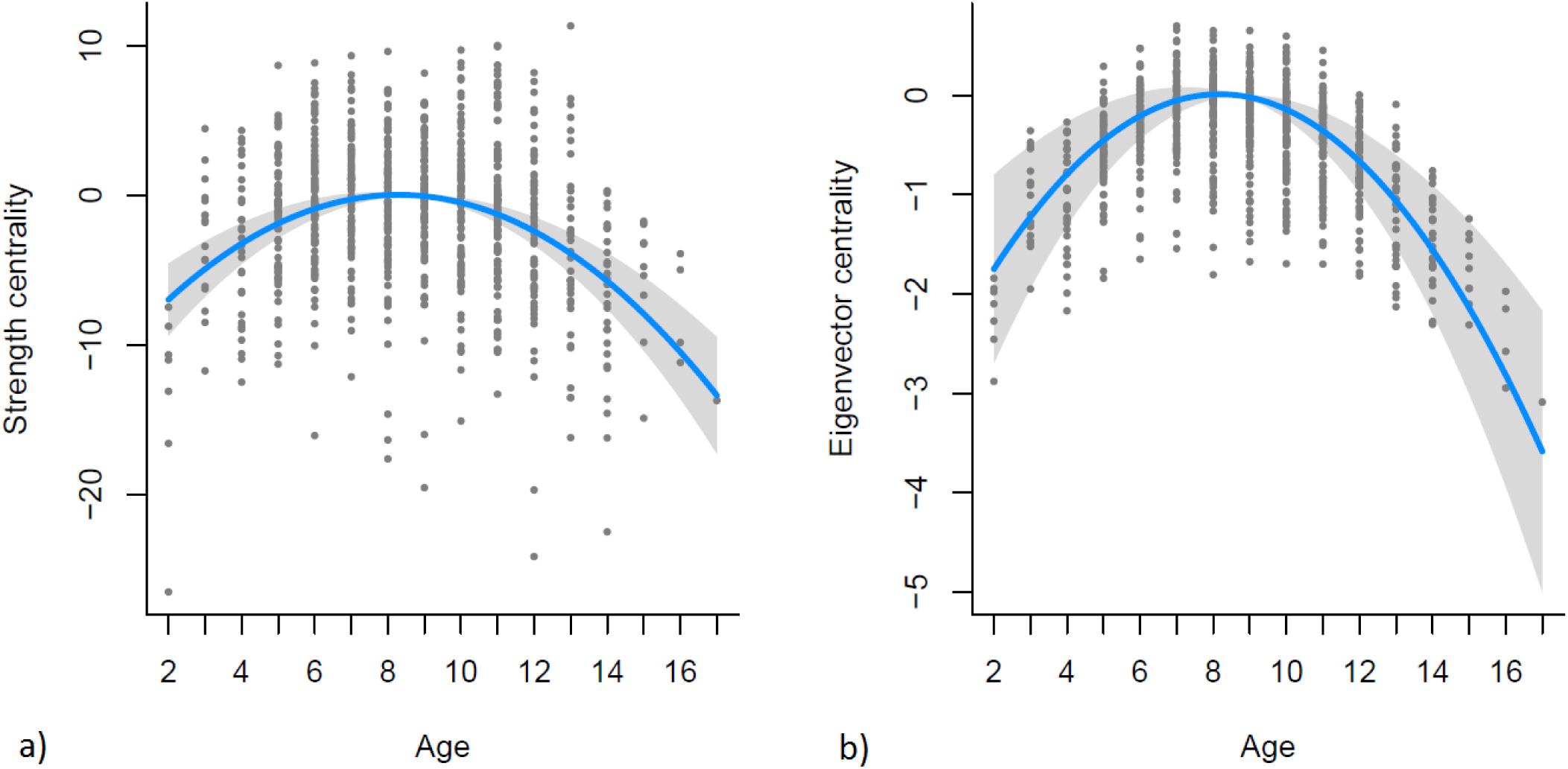
Partial regression plots representing the relationship between a) strength and b) eigenvector centrality respectively with age. Plots were built with the package visreg (Breheny & Burchett, 2017). The Y axis is scaled around the mean value of the response variable.

Consistently with results for strength centrality, eigenvector centrality was significantly lower in the summer season and was correlated with age, increasing until around 8-9 years and then decreasing when animals became older (table 5, figure 2). Eigenvector centrality was also negatively correlated with the occurrence of death in the months following the observations. ID was retained as a random factor in the selected model (ΔAIC between the selected model with and without ID as random factor was 13.5). The coefficients of determination of the selected model were: R^2^m=0.12, R^2^c=0.55.

### Seasonal and annual changes in the social structure

Annual and seasonal measures calculated to describe the global structure of the network are presented in table 6. The density of the network was close to one in all the years of the study with little variation (min.= 0.82, max.=0.99). TGS and gregariousness showed more variation over the years and between spring and summer seasons with both TGS (*β*=-6.66, Std. Err=2.05, *p*=0.005) and gregariousness (*β*=-5.09, Std. Err=1.43, *p*=0.002) being significantly lower during summer than during spring seasons (with no effect of sampling effort in either model). Gregariousness also showed differences in the coefficient of variation between years (min=0.18, max=0.40). Neither the proportion of old individuals nor the total number of males in the population had a significant effect on density or gregariousness. The proportion of old individuals also had no effect on TGS.

**Table 6.** Global measures calculated for the 2008-2017 networks. Density was calculated for the summer networks. Gregariousness (with coefficient of variation, CV, calculated as the ratio of the standard deviation to the mean) and TGS were calculated for both spring and summer networks.

| year | N tot | Proportion<br>ind > 9 y | Density | Gregariousness<br>spring mean (CV) | Gregariousness<br>summer mean (CV) | TGS<br>spring | TGS<br>summer |
| --- | --- | --- | --- | --- | --- | --- | --- |
| 2008 | 59 | 0.32 | 0.89 | 6.01 (0.20) | 3.09 (0.40) | 12.03 | 8.17 |
| 2009 | 55 | 0.31 | 0.82 | 5.68 (0.34) | 2.74 (0.32) | 14.72 | 11.65 |
| 2010 | 60 | 0.29 | 0.95 | 9.36 (0.26) | 6.85 (0.25) | 15.65 | 13.16 |
| 2011 | 57 | 0.26 | 0.92 | 7.70 (0.28) | 5.58 (0.36) | 11.90 | 10.08 |
| 2012 | 58 | 0.33 | 0.95 | 12.66 (0.22) | 7.59 (0.28) | 18.67 | 12.93 |
| 2013 | 61 | 0.43 | 0.89 | 12.26 (0.21) | 6.55 (0.31) | 20.50 | 9.90 |
| 2014 | 58 | 0.45 | 0.99 | 9.20 (0.20) | 6.76 (0.23) | 17.38 | 13.42 |
| 2015 | 63 | 0.40 | 0.98 | 16.51 (0.19) | 6.19 (0.27) | 27.07 | 11.27 |
| 2016 | 84 | 0.25 | 0.96 | 18.07 (0.18) | 9.20 (0.27) | 27.83 | 18.84 |
| 2017 | 84 | 0.19 | 0.99 | 11.56 (0.26) | 6.84 (0.29) | 22.65 | 16.71 |

## Discussion

We analysed the social network structure of male Alpine ibex using association and behavioural data collected over ten years. Male and female Alpine ibex are strongly segregated all year round, except for during the rutting season that occurs in December-January (Ruckstuhl and Neuhaus 2001). As they do not associate during spring and summer, our study did not consider interactions between males and females and focused on the social network structure of males alone.

### Association patterns

The association network for male Alpine ibex formed one discrete component, with a high density of connections. Fission-fusion dynamics lead almost every male in the population to associate with each other male at least once by being members of the same sub-group; field observations showed frequent splitting and merging events and that all males occasionally joined together in a single group. For example, during bad weather (i.e., mainly thunderstorms), male ibex tend to group at specific places used as shelters, usually at a lower altitude (A. Brambilla, *pers. obs*.). The Alpine ibex has a relatively small home range (Parrini et al., (2003) reported an average yearly home range of around 450-500 ha for males in the same study area) and lives in open high-altitude alpine habitat. During summer, males feed in large alpine pasturelands where resources are not concentrated in specific sites making resource attraction unlikely to explain the high network density observed in this species. The high density of connections observed in male Alpine ibex was therefore probably due to the general gregariousness of the species (Villaret & Bon, 1998) that is also observed in other mountain ungulates with a similar social structure and mating system (e.g., Bighorn sheep). The Alpine ibex is a polygynous species with males actively tending oestrus females and preventing subordinate males from approaching them (Willisch & Neuhaus, 2010; Willisch et al., 2012; Apollonio et al., 2013). As the rutting season of most mountain ungulates occurs in winter, when environmental conditions can be harsh and moving in the snow is energetically demanding (Signer et al., 2011), the establishment of the hierarchies often begins earlier. Indeed, agonistic interactions in Alpine ibex occur all year round except immediately after the rutting season (Brivio et al., 2010) and are more frequent during spring and summer months. Being part of a highly connected network may allow males of polygynous species living in highly seasonal environments to interact with many other males reducing the need of agonistic contests during winter, when it is more energetically costly. This advantage may compensate for the costs of group-living that, in the case of gregarious ungulates, are probably mostly related to infection risk (Brambilla et al., 2013; Marchand et al., 2017) as highly connected networks allows for potentially fast spreading of diseases (Wey et al., 2008; Marchand et al., 2017).

We found significant correlations between association matrices for spring and summer seasons within all the ten years of the study and also between adjacent years. The correlation coefficients were rather low, requiring caution in the interpretation of these results. However, the fact that all correlations between spring and summer association matrices of the same year and between summer association matrices of adjacent years were significant, seems to indicate that there are consistent associations within the population, both across seasons and years. These stable associations must convey some advantage as they are maintained despite a negligible risk of predation in this population and despite seasonal changes to the global association network structure. Our analyses carried out across a ten year period revealed that the factors leading to assortativity in Alpine ibex appeared constant throughout the study period. Males tended to associate with individuals of a similar age; dominance rank did not seem to predict associations, but difference in age did predict the likelihood of engaging in dominance interactions. The finding that age predicts association confirmed our expectation and supports the results of previous studies (*Capra ibex*, Villaret & Bon, 1995; *Capra pyrenaica*, Alados, 1986). However, we were not able to disentangle whether preferential associations were also present within individuals in the same age class and which factors drove these associations (e.g., kinship as observed by Cassinello & Calabuig (2008) in aoudades). A possible explanation of the fact that age predicts association is that individuals with similar characteristics (e.g., age and hence body size, Bergeron et al., 2010) also have similar energetic requirements and this leads them to associate and synchronize their activity so as to maintain group cohesion (Ruckstuhl & Neuhaus, 2000; Conradt & Roper, 2000). The large differences in body size of male Alpine ibex of different ages (Bergeron et al., 2010) further strengthen this explanation and in addition suggests that life history as well can contribute to the evolution of social structures. Furthermore, individuals may associate with others of similar age or size because they share similar social motivations and behaviours that enhance social cohesion and increase association (Cransac et al., 1998; Bon et al., 2001). In a polygynous species with male-male competition that lives in a strongly seasonal environment such as the Alps, adult males may share the need to interact to establish hierarchies before the reproductive season (Brivio et al., 2010; Willisch & Neuhaus, 2009). At the same time, young males may share the need to interact with other males through sparring to increase fighting skills, as observed for example in African elephants (Chiyo et al., 2011). In this Alpine ibex population, groups of males of similar age appear to merge and split with other groups regardless of their age. An implication of this could be that there is not a constant transfer of information between old and young males. In this species, the transfer of information between old and young individuals, if present, is likely to be more important for females that have to learn where to find safe and suitable places for parturition and the weaning of calves. In the case of males, since resources are homogeneously distributed during the summer season and predation risk is negligible, young individuals may avoid the costs of being constantly associated with mature males (with different time activity rhythm and physiological needs) but yet join them on some occasions.

As expected, and not surprisingly, age difference was also positively correlated with an increased likelihood of dominance interactions. Adult individuals were more frequently dominant, as has also been observed in other studies on the same species and on other polygynous mountain ungulates (Pelletier & Festa-Bianchet, 2006; Bergeron et al., 2010; Willisch et al., 2012; Apollonio et al., 2013). The Alpine ibex is a capital breeder with a slow life history strategy and reaches social sexual maturity at around 9-10 years of age (Willisch et al., 2012). Young individuals are therefore often in the lower positions of the hierarchy (Bergeron et al., 2010; Willisch et al., 2012). However, we did not find a correlation between likelihood of association and dominance rank differences, indicating that dominance rank is not a factor that seems to predict social preferences. Only a few of the agonistic interactions among males escalate into physical contact. When an individual’s dominance is clear, most of the interactions consists of threat displays. Therefore, associating with individuals of higher rank may not be costly, in terms of a risk of being injured, for low-rank males of species that evolved threat displays as a mean to establish hierarchies.

### Factors affecting node measures and seasonal network structure

Analysing the network at the node level, we also found that age predicted the position of the individuals in the network; as we were expecting, both strength centrality and eigenvector centrality showed a significant quadratic relationship with age. We found lower centrality values for young and old individuals and maximum centrality values for adult individuals at the peak of fitness (9-10 years old). Young individuals may join groups of adults but then leave and form separate groups, thus explaining the lower centrality values observed in this age class. Moreover, in accordance with the known life history of the species, old and senescent individuals also showed lower strength centrality and eigenvector centrality values. Because of their conservative life history strategy, adult survival is very high in Alpine ibex (ToÏgo et al., 2007). Many males therefore reach senescence with progressively deteriorating body condition (Bergeron et al., 2008; Brambilla et al., 2015) and changes in activity rhythm and spatial behaviour, meaning they are less likely to remain central in the network. Our results suggests that social behaviour changes in Alpine ibex may occur as a result of senescence (Siracusa et al., 2022). Similar patterns were recently observed in red deer where Albery et al., (2021a, preprint) found that older animals changed their spatial behaviour and became less socially connected. In accordance with our interpretation, we also found that strength and eigenvector centrality were lower if an animal died within subsequent months. Further evidence for this explanation came from field observations as well as from the structure of the network graphs (fig.1), where it appears that some individuals occupied a more peripheral position in the months preceding their death (fig.1, fig. S1). In particular, this seemed to happen for individuals that died from medium to long term diseases (e.g., pneumonia). In the months preceding death, those individuals were often observed alone, away from the other males. This likely happened because the progressive deterioration of their physical condition did not allow them to follow the daily movements and rhythm of the other individuals. It has been observed that Alpine ibex significantly decrease investment in horn growth in the years preceding death (von Hardenberg et al., 2004) suggesting that energetic investment in non-vital traits is reduced when the animal is in poor physical condition. Our observations suggest that the same is also true for behavioral traits. The observation of changes in individual behavior due to physical conditions and health status may have important implications for understanding the spreading dynamic of infectious diseases (Craft, 2015; Silk et al., 2019). However, we were not able to establish the cause of death for all the individuals and therefore our interpretation of this result remains speculative; a more detailed collection of data on the presence of infectious disease could help shed light on this important issue.

We also found preliminary evidence that inter-individual differences other than age are likely to play a role in determining social network positions in male ibex. Interestingly, an individual’s identity concurred to explain their association network position in terms of their strength centrality, suggesting that sociability varies consistently between individuals independent of their age. The presence of behavioural differences that are repeatable over time and across situations (i.e., personality or temperament) in animals has become widely recognized in the last few decades (Réale et al., 2007) and, since then, it has been observed in several species, including mountain ungulates (e.g., in bighorn sheep, Réale et al., 2000; Réale & Festa-Bianchet, 2003; Poissant et al., 2013). Sociability is one of the five temperament traits described in literature (Réale et al., 2007) and our ten-year dataset provided a unique opportunity to follow the associations of several individuals for many years, highlighting the potential impact of personality on network position. However, we have only found very preliminary evidence of repeatable individual differences in sociability (i.e., retention of the individual identity as a random effect in the selected model) while other measurements of sociability as well as its heritability should be assessed in order to reliably quantify personality in this species (Réale et al., 2010).

### Seasonal and annual changes in the social structure

Analysing seasonal differences in the social structure, we found that mean strength and mean eigenvector centrality in the association network, gregariousness and Typical Group Size (TGS) were all significantly lower in the summer compared with the spring. The overall higher values of sociality observed during spring compared to summer can partly be explained by seasonal differences in resource availability (Peignier et al., 2019). During spring, resources are more concentrated as there are only a few pastures available at low altitude, where snow melts earlier. Hence, all male Alpine ibex group together there following the winter. In summer, as snow melts and more pastures become available, male Alpine ibex migrate to a higher altitude (Parrini et al., 2003) and therefore have fewer constraints on where to feed and with whom to associate. Accordingly, overall gregariousness and TGS decrease in addition to the mean strength centrality. Similar results have been found in other ungulates, for example in reindeer (*Rangifer tarandus*): Peignier et al., (2019) observed low home range overlap and social associations in summer, when resources were distributed homogeneously, but higher home range overlap and social associations in winter, when resources were distributed heterogeneously. Our results hence strengthen the hypothesis that ecological constrains may influence social behaviour in ungulates. On the other side, in horses, strength centrality was found to be lower outside of the mating season; this highlighted a similar increased freedom in the choice of social partners and feeding locations when constraints were reduced, but in this species, the constraints appeared to be social rather than ecological (Stanley et al., 2018).

Finally, we aimed to verify if the overall structure of the social network remained consistent over a period of ten years and whether possible changes could be explained by demographic characteristics of the population. Due to their experience and knowledge, older individuals are expected to help maintain group cohesion, e.g., leading group movement (McComb et al., 2001). Despite most of the study on this topic being conducted on females in matrilineal societies (McComb et al., 2001; Brent et al., 2015), it has recently been shown that the same may apply also to males (Allen et al., 2020). We therefore expected to find the highest values for global centrality measures in years with a higher number of mature individuals. However, the proportion of males over 9 years of age seemed not to affect any of the global measures considered (TGS, gregariousness, density). Also the total number of individuals in the population did not affect either gregariousness or density. The absence of a relationship between global network measures and the demographic structure of the population could be due to the seasonal movement patterns apparent in this species. Unlike animals such as African elephants, the home range of the Alpine ibex in our study area is rather small, around 450-500 ha (Parrini et al., 2003). The seasonal movement of most male Alpine ibex occurs within the same valley, and they show a high site fidelity, possibly because they live in a rather predictable environment (Morrison et al., 2021). This likely makes the role of old, experienced individuals less crucial. Similar conditions may apply to other mountain ungulates with low dispersal rate and high site fidelity (Festa-Bianchet, 1986). If this hypothesis is confirmed, it could mean that the social cohesion of male mountain ungulates group is stable and is not dependent on the presence of old and experienced individuals, i.e., it shows resilience to changes in group membership. However, we have to acknowledge that, during the years of the study, the demographic structure of the population was not subject to drastic changes as the environmental conditions remained relatively stable and some old males were alive in the population across all the years of the study, even after harsh winters. This could mean that the presence of a few older individuals may be enough to maintain social cohesion; however, it could also signify that we did not capture enough variation to draw biologically meaningful conclusions. As this aspect could have important conservation implications, further data should be collected to determine the effect of demographic structure on the social structure of mountain ungulates. Particularly, building networks on a finer scale (e.g., daily networks) coupled with more detailed data collection on associations (e.g., using the nearest neighbour method or conducting focal observations), as well as accounting for spatial behaviour (Albery et al., 2021b), could help in better understanding the factors shaping associations and the global network structure.

In conclusion, this study investigated social network dynamics of male of a mountain ungulate over a relatively long time period focusing at the level of group and individual, allowing us to explore ungulate sociality over different scales. As most previous studies on ungulate social networks have focused on female associations, this is one of the few to quantify male social structure in a gregarious ungulate with high levels of sexual segregation. We found that the male Alpine ibex social network is highly cohesive with all individuals being connected; this has important implications for the management of disease outbreaks. Social structure changed during seasons, suggesting that ecological constrains such as resource availability may play a role in shaping associations in gregarious ungulates. However, we also demonstrated that male Alpine ibex did not associate randomly but showed stable associations, and that the strength of these associations varies across individuals, with age similarity being a factor driving these associations. This, in turn, suggests that ecological constrains are not enough to explain gregariousness and preferential associations in ungulates; these are likely to also be driven by physiological and social needs, ultimately varying according to individual characteristics and species life history. Finally, our results highlight the importance of long-term studies based on individually recognizable subjects (Festa-Bianchet et al., 2017).

## Acknowledgements

Version 3 of this preprint has been peer-reviewed and recommended by Peer Community In Network Science. https://doi.org/10.24072/pci.networksci.100004 We thank Brenda McCowan and Sandra Smith Aguilar for constructive reviews of the manuscript and Gabriel Ramos-Fernández for the recommendation and the evaluation process.

We deeply thank Bruno Bassano from the Gran Paradiso National Park biodiversity and scientific research service and the Park Rangers of the surveillance service for captures and marking of Alpine ibex and for providing logistic support during field work. We are grateful to all the students and researchers that helped with data collection in the ten years of the study and to Edward Monovich for the artwork. We also thank David Laniado and Cristian Pasquaretta for helpful discussion and suggestion on data analysis.

## Data, scripts and codes availability

The data used to produce this manuscript are accessible on the Dryad Digital Repository with the following DOI: https://doi.org/10.5061/dryad.w0vt4b8st

The code used for the data analysis contained in this manuscript is accessible at Zenodo with the following DOI: https://doi.org/10.5281/zenodo.6499633

## Supplementary material

Supplementary information S0, S1 and S2 are deposited on bioRxiv as Supplementary Material of the manuscript with the following DOI https://doi.org/10.1101/2021.12.02.470954

### Conflict of interest disclosure

The authors declare they have no conflict of interest relating to the content of this article. Cédric Sueur is a recommender for PCI Network Science.

### Funding

The authors conducted this research as part of their academic activity. This research did not receive specific grant from funding agencies.

Supplementary information S0, S1 and S2 are deposited on bioRxiv as Supplementary Material of the manuscript with the following DOI https://doi.org/10.1101/2021.12.02.470954

## Appendix

**Supplementary Information S0** Number of days of observation, number of surveys and of group observed, number of unique individuals and total number of animals observed during spring and summer season 2008-2017

**Supplementary Information S1** graphical representation of the weighted association networks of male Alpine ibex in spring and summer season of each year of the study (2008-2018).

**Supplementary Information S2** Model selection for the GLMM performed to explain the variance of Strength Centrality Eand igenvector Centrality

## Notes

https://doi.org/10.5061/dryad.w0vt4b8st

https://doi.org/10.5281/zenodo.6499633

### Competing Interest Statement

The authors have declared no competing interest.

## References

Akaike, H. (1973). Information theory and the maximum likelihood principle in 2nd International Symposium on Information Theory (B.N. Petrov and F. Csäki, eds.). Akademiai Kiàdo, Budapest.

Alados, C. L. (1986). Spatial structure in groups of Spanish ibex (Capra pyrenaica). Biol Behav, 11, 176–185.

Albers, P. C., & Vries, H. D. (2001). Elo-rating as a tool in the sequential estimation of dominance strengths. Animal Behaviour, 489–495. https://doi.org/10.1006/anbe.2000.1571

Albery, G. F., Clutton-Brock, T. H., Morris, A., Morris, S., Pemberton, J. M., Nussey, D. H., & Firth, J. A. (2021a). Ageing red deer alter their spatial behaviour and become less social. bioRxiv. https://doi.org/10.1101/2021.06.11.448092

Albery, G. F., Morris, A., Morris, S., Pemberton, J. M., Clutton-Brock, T. H., Nussey, D. H., & Firth, J. A. (2021b). Multiple spatial behaviours govern social network positions in a wild ungulate. Ecology Letters, 24(4), 676–686. https://doi.org/10.1111/ele.13684

Allen, C. R., Brent, L. J., Motsentwa, T., Weiss, M. N., & Croft, D. P. (2020). Importance of old bulls: leaders and followers in collective movements of all-male groups in African savannah elephants (Loxodonta africana). Scientific reports, 10(1), 1–9. https://doi.org/10.1038/s41598-020-70682-y

Altmann, J. (1974). Observational study of behavior: sampling methods. Behaviour, 49(3-4), 227–266. https://doi.org/10.1163/156853974X00534

Apollonio, M., Brivio, F., Rossi, I., Bassano, B., & Grignolio, S. (2013). Consequences of snowy winters on male mating strategies and reproduction in a mountain ungulate. Behavioural processes, 98, 44–50. https://doi.org/10.1016/j.beproc.2013.05.001

Appleby M. C. (1983). Competition in a red deer stag social group: rank, age and relatedness of opponents. Animal Behaviour, 31, 913–918. https://doi.org/10.1016/S0003-3472(83)80246-2

Aureli F. et al. 2008. Fission–fusion dynamics: new research frameworks. Current Anthropology, 49: 627–654. https://doi.org/10.1086/586708

Bassano, B., von Hardenberg, A., Pelletier, F., & Gobbi, G. (2003). A method to weigh free-ranging ungulates without handling. Wildlife Society Bulletin, 1205–1209. https://www.jstor.org/stable/3784468

Bastian, M., Heymann, S., & Jacomy, M. (2009, March). Gephi: an open source software for exploring and manipulating networks. In Proceedings of the international AAAI conference on web and social media (Vol. 3, No. 1, pp. 361–362).

Barton, K. (2009) Mu-MIn: Multi-model inference. R Package Version 1.43.17 https://CRAN.R-project.org/package=MuMIn

Bates D, Mächler M, Bolker B & Walker S (2015). Fitting Linear Mixed-Effects Models Using lme4. Journal of Statistical Software, 67(1), 1–48. https://doi.org/10.18637/jss.v067.i01

Bergeron, P., Festa-Bianchet, M., Von Hardenberg, A., & Bassano, B. (2008). Heterogeneity in male horn growth and longevity in a highly sexually dimorphic ungulate. Oikos, 117(1), 77–82. https://doi.org/10.1111/j.2007.0030-1299.16158.x

Bergeron, P., Grignolio, S., Apollonio, M., Shipley, B., & Festa-Bianchet, M. (2010). Secondary sexual characters signal fighting ability and determine social rank in Alpine ibex (Capra ibex). Behavioral Ecology and Sociobiology, 64(8), 1299–1307. https://doi.org/10.1007/s00265-010-0944-x

Bon, R., Rideau, C., Villaret, J. C., & Joachim, J. (2001). Segregation is not only a matter of sex in Alpine ibex, Capra ibex ibex. Animal Behaviour, 62(3), 495–504. https://doi.org/10.1006/anbe.2001.1776

Bond, M. L., Lee, D. E., Ozgul, A., & König, B. (2019). Fission–fusion dynamics of a megaherbivore are driven by ecological, anthropogenic, temporal, and social factors. Oecologia, 191(2), 335–347. https://doi.org/10.1007/s00442-019-04485-y

Borgeaud, C., Sosa, S., Sueur, C., & Bshary, R. (2017). The influence of demographic variation on social network stability in wild vervet monkeys. Animal Behaviour, 134, 155–165. https://doi.org/10.1016/j.anbehav.2017.09.028

Brambilla, A., von Hardenberg, A., Kristo, O., Bassano, B., & Bogliani, G. (2013). Don’t spit in the soup: faecal avoidance in foraging wild Alpine ibex, Capra ibex. Animal Behaviour, 86(1), 153–158. https://doi.org/10.1016/j.anbehav.2013.05.006

Brambilla, A., & Brivio, F. (2018). Assessing the effects of helicopter disturbance in a mountain ungulate on different time scales. Mammalian Biology, 90(1), 30–37. https://doi.org/10.1016/j.mambio.2018.02.001

Brambilla, A., & Canedoli, C. (2014). How to continue measuring horn growth after capture in Alpine ibex. Journal of Mountain Ecology, 9.

Brambilla, A., Biebach, I., Bassano, B., Bogliani, G., & von Hardenberg, A. (2015). Direct and indirect causal effects of heterozygosity on fitness-related traits in Alpine ibex. Proceedings of the Royal Society B: Biological Sciences, 282(1798), 20141873. https://doi.org/10.1098/rspb.2014.1873

Brambilla, A., Keller, L., Bassano, B., & Grossen, C. (2018). Heterozygosity–fitness correlation at the major histocompatibility complex despite low variation in Alpine ibex (Capra ibex). Evolutionary applications, 11(5), 631–644. https://doi.org/10.1111/eva.12575

Brambilla, A., Von Hardenberg, A., Nelli, L., & Bassano, B. (2020). Distribution, status, and recent population dynamics of Alpine ibex Capra ibex in Europe. Mammal Review, 50(3), 267–277. https://doi.org/10.1111/mam.12194

Breheny, P and Burchett, W (2017). Visualization of Regression Models Using visreg. The R Journal, 9: 56–71. https://doi.org/10.32614/RJ-2017-046

Brent, L. J., Lehmann, J., & Ramos-Fernández, G. (2011). Social network analysis in the study of nonhuman primates: A historical perspective. American Journal of Primatology, 73(8), 720–730. https://doi.org/10.1002/ajp.20949

Brent, L. J., Franks, D. W., Foster, E. A., Balcomb, K. C., Cant, M. A., & Croft, D. P. (2015). Ecological knowledge, leadership, and the evolution of menopause in killer whales. Current Biology, 25(6), 746–750. https://doi.org/10.1016/j.cub.2015.01.037

Brivio, F., Grignolio, S., & Apollonio, M. (2010). To feed or not to feed? Testing different hypotheses on rut-induced hypophagia in a mountain ungulate. Ethology, 116(5), 406–415. https://doi.org/10.1111/j.1439-0310.2010.01753.x

Brivio, F., Grignolio, S., Sica, N., Cerise, S., & Bassano, B. (2015). Assessing the impact of capture on wild animals: the case study of chemical immobilisation on alpine ibex. PLoS One, 10(6), e0130957. https://doi.org/10.1371/journal.pone.0130957

Burnham, K. P., Anderson, D. R., & Huyvaert, K. P. (2011). AIC model selection and multimodel inference in behavioral ecology: some background, observations, and comparisons. Behavioral ecology and sociobiology, 65(1), 23–35. https://doi.org/10.1007/s00265-010-1029-6

Butts, C. T., (2020). sna: Tools for Social Network Analysis. R package version 2.6. https://CRAN.R-project.org/package=sna

Cameron, E. Z., Setsaas, T. H., & Linklater, W. L. (2009). Social bonds between unrelated females increase reproductive success in feral horses. Proceedings of the National Academy of Sciences, 106(33), 13850–13853. https://doi.org/10.1073/pnas.0900639106

Cantor, M., Wedekin, L. L., Guimaraes, P. R., Daura-Jorge, F. G., Rossi-Santos, M. R., & Simoes-Lopes, P. C. (2012). Disentangling social networks from spatiotemporal dynamics: the temporal structure of a dolphin society. Animal Behaviour, 84(3), 641–651. https://doi.org/10.1016/j.anbehav.2012.06.019

Carter, K. D., Brand, R., Carter, J. K., Shorrocks, B., & Goldizen, A. W. (2013). Social networks, long-term associations and age-related sociability of wild giraffes. Animal Behaviour, 86(5), 901–910. https://doi.org/10.1016/j.anbehav.2013.08.002

Cassinello, J. and Calabuig, G. (2008). Spatial Association in a Highly Inbred Ungulate Population: Evidence of Fine-Scale Kin Recognition. Ethology, 114: 124–132. https://doi.org/10.1111/j.1439-0310.2007.01453.x

Chiyo, P. I., Archie, E. A., Hollister-Smith, J. A., Lee, P. C., Poole, J. H., Moss, C. J., & Alberts, S. C. (2011). Association patterns of African elephants in all-male groups: the role of age and genetic relatedness. Animal Behaviour, 81(6), 1093–1099. https://doi.org/10.1016/j.anbehav.2011.02.013

Cinar, O., & Viechtbauer, W. (2021). poolr: Methods for Pooling P-Values from (Dependent) Tests. R package version 1.0-0. https://CRAN.R-project.org/package=poolr

Clutton-Brock, T. (2021). Social evolution in mammals. Science, 373(6561), eabc9699. https://doi.org/10.1126/science.abc9699

Connor, R. C., & Wells, R. S. (2000). The bottlenose dolphin: social relationships in a fission–fusion society. In ‘Cetacean Societies: Field Studies of Dolphins and Whales’. Eds J. Mann, RC Connor, PL Tyack and H. Whitehead. pp. 91–126.

Conradt, L., & Roper, T. J. (2000). Activity synchrony and social cohesion: a fission-fusion model. Proceedings of the Royal Society of London. Series B: Biological Sciences, 267(1458), 2213–2218. https://doi.org/10.1098/rspb.2000.1271

Couturier M.A.J. (1962) Le Bouquetin des Alps: Capra aegagrus ibex ibex. L. Arthaud, Grenoble France

Couzin, I. D., & Laidre, M. E. (2009). Fission–fusion populations. Current biology, 19(15), R633–R635. https://doi.org/10.1016/j.cub.2009.05.034

Cransac, N., Gerard, J. F., Maublanc, M. L., & Pépin, D. (1998). An example of segregation between age and sex classes only weakly related to habitat use in mouflon sheep (Ovis gmelini). Journal of Zoology, 244(3), 371–378. https://doi.org/10.1111/j.1469-7998.1998.tb00042.x

Craft, M. E. (2015). Infectious disease transmission and contact networks in wildlife and livestock. Philosophical Transactions of the Royal Society B: Biological Sciences, 370(1669), 20140107. https://doi.org/10.1098/rstb.2014.0107

Croft D. P., James R., Krause J. (2008). Exploring Animal Social Networks. Princeton University Press. https://doi.org/10.1515/9781400837762

Elo A. E. (1978). The Rating of Chess Players, Past and Present. New York: Arco.

Festa-Bianchet, M. (1986). Site fidelity and seasonal range use by bighorn rams. Canadian Journal of Zoology, 64(10), 2126–2132. https://doi.org/10.1139/z86-326

Festa-Bianchet, M., Douhard, M., Gaillard, J. M., & Pelletier, F. (2017). Successes and challenges of long-term field studies of marked ungulates. Journal of Mammalogy, 98(3), 612–620. https://doi.org/10.1093/jmammal/gyw227

Firth, J. A., Cole, E. F., Ioannou, C. C., Quinn, J. L., Aplin, L. M., Culina, A., … & Sheldon, B. C. (2018). Personality shapes pair bonding in a wild bird social system. Nature ecology & evolution, 2(11), 1696–1699. https://doi.org/10.1038/s41559-018-0670-8

Fischhoff, I. R., Dushoff, J., Sundaresan, S. R., Cordingley, J. E., & Rubenstein, D. I. (2009). Reproductive status influences group size and persistence of bonds in male plains zebra (Equus burchelli). Behavioral Ecology and Sociobiology, 63(7), 1035–1043. https://doi.org/10.1007/s00265-009-0723-8

Flack, J. C., Girvan, M., De Waal, F. B., & Krakauer, D. C. (2006). Policing stabilizes construction of social niches in primates. Nature, 439(7075), 426–429. https://doi.org/10.1038/nature04326

Fogarty S., Cote J., Sih A. 2011. Social personality polymorphism and the spread of invasive species: a model. The American Naturalist, 177(3), 273–287. https://doi.org/10.1086/658174

Franks, D. W., Ruxton, G. D., & James, R. (2010). Sampling animal association networks with the gambit of the group. Behavioral ecology and sociobiology, 64(3), 493–503. https://doi.org/10.1007/s00265-009-0865-8

Godde, S., Humbert, L., Côté, S. D., Réale, D., & Whitehead, H. (2013). Correcting for the impact of gregariousness in social network analyses. Animal Behaviour, 85(3), 553–558. https://doi.org/10.1016/j.anbehav.2012.12.010

Grueter, C. C., Qi, X., Zinner, D., Bergman, T., Li, M., Xiang, Z., … & Swedell, L. (2020). Multilevel organisation of animal sociality. Trends in ecology & evolution. 35(9), 834–847. https://doi.org/10.1016/j.tree.2020.05.003

Handcock, M. S., Hunter, D. R., Butts, C. T., Goodreau, S. M., & Morris, M. (2008). statnet: Software tools for the representation, visualization, analysis and simulation of network data. Journal of Statistical Software, 24(1), 1548. https://doi.org/10.18637/jss.v024.i01

Holekamp, K. E., Cooper, S. M., Katona, C. I., Berry, N. A., Frank, L. G., & Smale, L. (1997). Patterns of association among female spotted hyenas (Crocuta crocuta). Journal of Mammalogy, 78(1), 55–64. https://doi.org/10.2307/1382638

Holm, S. (1979). A simple sequentially rejective multiple test procedure. Scandinavian journal of statistics, 65–70.

Ilany, A., & Akcay, E. (2016). Social inheritance can explain the structure of animal social networks. Nature communications, 7(1), 1–10. https://doi.org/10.1038/ncomms12084

Jarman, P. (1974). The social organisation of antelope in relation to their ecology. Behaviour, 48(1-4), 215-267. https://doi.org/10.1163/156853974X00345

Jewell P. A., Milner C., Boyd J. M. (1974). Island survivors. The ecology of the Soay sheep of St. Kilda. The Athlone Press, University of London, London

Kimura, R. (1998). Mutual grooming and preferred associate relationships in a band of free-ranging horses. Applied Animal Behaviour Science, 59(4), 265–276. https://doi.org/10.1016/S0168-1591(97)00129-9

Kerth, G., & Konig, B. (1999). Fission, fusion and nonrandom associations in female Bechstein’s bats (Myotis bechsteinii). Behaviour, 136(9), 1187–1202. https://doi.org/10.1163/156853999501711

Krackhardt, D. (1988). Predicting with networks: Nonparametric multiple regression analysis of dyadic data. Social networks, 10(4), 359–381. https://doi.org/10.1016/0378-8733(88)90004-4

Krause, J., Ruxton, G. D., Ruxton, G., & Ruxton, I. G. (2002). Living in groups. Oxford University Press.

Krause, J., Lusseau, D., & James, R. (2009). Animal social networks: an introduction. Behavioral Ecology and Sociobiology, 63(7), 967–973. https://doi.org/10.1007/s00265-009-0747-0

Krause, J., James, R., & Croft, D. P. (2010). Personality in the context of social networks. Philosophical Transactions of the Royal Society B: Biological Sciences, 365(1560), 4099–4106. https://doi.org/10.1098/rstb.2010.0216

Krause, J., James, R., Franks, D. W., & Croft, D. P. (Eds.). (2015). Animal social networks. Oxford University Press, USA. https://doi.org/10.1093/acprof:oso/9780199679041.001.0001

Le Pendu, Y., Briedermann, L., Gerard, J. F., & Maublanc, M. L. (1995). Inter-individual associations and social structure of a mouflon population (Ovis orientalis musimon). Behavioural processes, 34(1), 67–80. https://doi.org/10.1016/0376-6357(94)00055-L

Machanda, Z. P., & Rosati, A. G. (2020). Shifting sociality during primate ageing. Philosophical Transactions of the Royal Society B, 375(1811), 20190620. https://doi.org/10.1098/rstb.2019.0620

MacIntosh, A. J., Jacobs, A., Garcia, C., Shimizu, K., Mouri, K., Huffman, M. A., & Hernandez, A. D. (2012). Monkeys in the middle: parasite transmission through the social network of a wild primate. PloS one, 7(12), e51144. https://doi.org/10.1371/journal.pone.0051144

Marchand, P., Freycon, P., Herbaux, J. P., Game, Y., ToÏgo, C., Gilot-Fromont, E., … & Hars, J. (2017). Sociospatial structure explains marked variation in brucellosis seroprevalence in an Alpine ibex population. Scientific reports, 7(1), 1–12. https://doi.org/10.1038/s41598-017-15803-w

McComb, K., Moss, C., Durant, S. M., Baker, L., & Sayialel, S. (2001). Matriarchs as repositories of social knowledge in African elephants. Science, 292(5516), 491–494. https://doi.org/10.1126/science.1057895

Morrison, T. A., Merkle, J. A., Hopcraft, J. G. C., Aikens, E. O., Beck, J. L., Boone, R. B., … & Kauffman, M. J. (2021). Drivers of site fidelity in ungulates. Journal of animal ecology, 90(4), 955–966. https://doi.org/10.1111/1365-2656.13425

Moscovice, L. R., Sueur, C., & Aureli, F. (2020). How socio-ecological factors influence the differentiation of social relationships: an integrated conceptual framework. Biology Letters, 16(9), 20200384. https://doi.org/10.1098/rsbl.2020.0384

Neumann, C., Duboscq, J., Dubuc, C., Ginting, A., Irwan, A. M., Agil, M., … & Engelhardt, A. (2011). Assessing dominance hierarchies: validation and advantages of progressive evaluation with Elo-rating. Animal Behaviour, 82(4), 911–921. https://doi.org/10.1016/j.anbehav.2011.07.016

Opsahl, T., 2009. Structure and Evolution of Weighted Networks. University of London (Queen Mary College), London, UK, pp. 104–122. http://toreopsahl.com/tnet/

Papageorgiou, D., & Farine, D. R. (2020). Multilevel societies in birds. Trends in Ecology & Evolution, 36. https://doi.org/10.1016/j.tree.2020.10.008

Parrini, F., Grignolio, S., Luccarini, S., Bassano, B., & Apollonio, M. (2003). Spatial behaviour of adult male Alpine ibex Capra ibex ibex in the Gran Paradiso National Park, Italy. Acta Theriologica, 48(3), 411–423. https://doi.org/10.1007/BF03194179

Peignier, M., Webber, Q. M., Koen, E. L., Laforge, M. P., Robitaille, A. L., & Vander Wal, E. (2019). Space use and social association in a gregarious ungulate: Testing the conspecific attraction and resource dispersion hypotheses. Ecology and evolution, 9(9), 5133–5145. https://doi.org/10.1002/ece3.5071

Pelletier, F., & Festa-Bianchet, M. (2006). Sexual selection and social rank in bighorn rams. Animal Behaviour, 71(3), 649–655. https://doi.org/10.1016/j.anbehav.2005.07.008

Pinter-Wollman, N., Hobson, E. A., Smith, J. E., Edelman, A. J., Shizuka, D., De Silva, S., … & Fewell, J. (2014). The dynamics of animal social networks: analytical, conceptual, and theoretical advances. Behavioral Ecology, 25(2), 242–255. https://doi.org/10.1093/beheco/art047

Poissant, J., Réale, D., Martin, J. G. A., Festa-Bianchet, M., & Coltman, D. W. (2013). A quantitative trait locus analysis of personality in wild bighorn sheep. Ecology and evolution, 3(3), 474–481. https://doi.org/10.1002/ece3.468

Podgórski, T., Lusseau, D., Scandura, M., Sönnichsen, L., & Jędrzejewska, B. (2014). Long-lasting, kin-directed female interactions in a spatially structured wild boar social network. PLoS One, 9(6), e99875. https://doi.org/10.1371/journal.pone.0099875

R Core Team (2020). R: A language and environment for statistical computing. Vienna, Austria: R Computing Foundation for Science. http://www.R-project.org

Ramos, A., Manizan, L., Rodriguez, E., Kemp, Y. J., & Sueur, C. (2019). The social network structure of a semi-free roaming European bison herd (Bison bonasus). Behavioural processes, 158, 97–105. https://doi.org/10.1016/j.beproc.2018.11.005

Réale, D., Dingemanse, N. J., Kazem, A. J., & Wright, J. (2010). Evolutionary and ecological approaches to the study of personality. Philosophical Transactions of the Royal Society B: Biological Sciences, 365(1560), 3937–3946. https://doi.org/10.1098/rstb.2010.0222

Réale, D., Gallant, B. Y., Leblanc, M., & Festa-Bianchet, M. (2000). Consistency of temperament in bighorn ewes and correlates with behaviour and life history. Animal behaviour, 60(5), 589–597. https://doi.org/10.1006/anbe.2000.1530

Réale, D., & Festa-Bianchet, M. (2003). Predator-induced natural selection on temperament in bighorn ewes. Animal behaviour, 65(3), 463–470. https://doi.org/10.1006/anbe.2003.2100

Réale, D., Reader, S. M., Sol, D., McDougall, P. T., & Dingemanse, N. J. (2007). Integrating animal temperament within ecology and evolution. Biological reviews, 82(2), 291–318. https://doi.org/10.1111/j.1469-185X.2007.00010.x

Ruckstuhl, K., & Neuhaus, P. (2001). Behavioral synchrony in ibex groups: effects of age, sex and habitat. Behaviour, 138(8), 1033–1046. https://doi.org/10.1163/156853901753286551

Ruckstuhl, K., & Neuhaus, P. (2000). Sexual segregation in ungulates: a new approach. Behaviour, 137(3), 361–377. https://doi.org/10.1163/156853900502123

Schakner, Z. A., Petelle, M. B., Tennis, M. J., Van der Leeuw, B. K., Stansell, R. T., & Blumstein, D. T. (2017). Social associations between California sea lions influence the use of a novel foraging ground. Royal Society open science, 4(5), 160820. https://doi.org/10.1098/rsos.160820

Siracusa, E. R., Higham, J. P., Snyder-Mackler, N., & Brent, L. J. (2022). Social ageing: exploring the drivers of late-life changes in social behaviour in mammals. Biology Letters, 18(3), 20210643. https://doi.org/10.1098/rsbl.2021.0643

Shizuka, D., & Johnson, A. E. (2020). How demographic processes shape animal social networks. Behavioral Ecology, 31(1), 1–11. https://doi.org/10.1093/beheco/arz083

Signer, C., Ruf, T., & Arnold, W. (2011). Hypometabolism and basking: the strategies of Alpine ibex to endure harsh over-wintering conditions. Functional Ecology, 25(3), 537–547. https://doi.org/10.1111/j.1365-2435.2010.01806.x

Silk, M. J., Croft, D. P., Tregenza, T., & Bearhop, S. (2014). The importance of fission–fusion social group dynamics in birds. Ibis, 156(4), 701–715. https://doi.org/10.1111/ibi.12191

Silk, M. J., Hodgson, D. J., Rozins, C., Croft, D. P., Delahay, R. J., Boots, M., & McDonald, R. A. (2019). Integrating social behaviour, demography and disease dynamics in network models: applications to disease management in declining wildlife populations. Philosophical Transactions of the Royal Society B, 374(1781), 20180211. https://doi.org/10.1098/rstb.2018.0211

Smith, J. E., Powning, K. S., Dawes, S. E., Estrada, J. R., Hopper, A. L., Piotrowski, S. L., & Holekamp, K.E. (2011). Greetings promote cooperation and reinforce social bonds among spotted hyaenas. Animal Behaviour, 81(2), 401–415. https://doi.org/10.1016/j.anbehav.2010.11.007

Smuts B. B., Cheney D. L., Seyfarth R. M., Wrangham R. W., Struhsaker T. T. (1987). Primate societies. Chicago: The University of Chicago Press. https://doi.org/10.7208/chicago/9780226220468.001.0001

Snijders, L., Blumstein, D. T., Stanley, C. R., & Franks, D. W. (2017). Animal social network theory can help wildlife conservation. Trends in ecology & evolution, 32(8), 567–577. https://doi.org/10.1016/j.tree.2017.05.005

Sosa, S. (2016). The influence of gender, age, matriline and hierarchical rank on individual social position, role and interactional patterns in Macaca sylvanus at ‘La Forêt des singes’: A multilevel social network approach. Frontiers in psychology, 7, 529. https://doi.org/10.3389/fpsyg.2016.00529

Sosa, S., Puga-Gonzalez, I., Hu, F., Pansanel, J., Xie, X., & Sueur, C. (2020). A multilevel statistical toolkit to study animal social networks: the Animal network toolkit Software (Ants) R package. Scientific reports, 10(1), 1–8. https://doi.org/10.1038/s41598-020-69265-8

Sosa, S., Sueur, C., & Puga-Gonzalez, I. (2021a). Network measures in animal social network analysis: Their strengths, limits, interpretations and uses. Methods in Ecology and Evolution, 12(1), 10–21. https://doi.org/10.1111/2041-210X.13366

Sosa, S., Jacoby, D. M., Lihoreau, M., & Sueur, C. (2021b). Animal social networks: Towards an integrative framework embedding social interactions, space and time. Methods in Ecology and Evolution, 12,4–9. https://doi.org/10.1111/2041-210X.13539

Stanley, C. R., & Dunbar, R. I. M. (2013). Consistent social structure and optimal clique size revealed by social network analysis of feral goats, Capra hircus. Animal Behaviour, 85(4), 771–779. https://doi.org/10.1016/j.anbehav.2013.01.020

Stanley, C. R., Mettke-Hofmann, C., Hager, R., & Shultz, S. (2018). Social stability in semiferal ponies: networks show interannual stability alongside seasonal flexibility. Animal Behaviour, 136, 175–184. https://doi.org/10.1016/j.anbehav.2017.04.013

Statnet Development Team (2003-2020). Statnet: Software tools for the Statistical Modeling of Network Data. http://statnet.org

Sueur, C., King, A. J., Conradt, L., Kerth, G., Lusseau, D., Mettke-Hofmann, C., … & Aureli, F. (2011). Collective decision-making and fission–fusion dynamics: a conceptual framework. Oikos, 120(11), 1608–1617. https://doi.org/10.1111/j.1600-0706.2011.19685.x

Tarlow, E. M., & Blumstein, D. T. (2007). Evaluating methods to quantify anthropogenic stressors on wild animals. Applied Animal Behaviour Science, 102(3-4), 429–451. https://doi.org/10.1016/j.applanim.2006.05.040

Teitelbaum, C. S., Converse, S. J., & Mueller, T. (2017). Birds choose long-term partners years before breeding. Animal Behaviour, 134, 147–154. https://doi.org/10.1016/j.anbehav.2017.10.015

ToÏgo, C., Gaillard, J. M., Festa-Bianchet, M.., Largo, E., Michallet, J., & Maillard, D. (2007). Sex-and age-specific survival of the highly dimorphic Alpine ibex: evidence for a conservative life-history tactic. Journal of Animal Ecology, 76(4), 679–686. https://doi.org/10.1111/j.1365-2656.2007.01254.x

Turner, J. W., Bills, P. S., & Holekamp, K. E. (2018). Ontogenetic change in determinants of social network position in the spotted hyena. Behavioral ecology and sociobiology, 72(1), 10. https://doi.org/10.1007/s00265-017-2426-x

van de Waal, E., & Bshary, R. (2011). Social-learning abilities of wild vervet monkeys in a two-step task artificial fruit experiment. Animal Behaviour, 81(2), 433–438. https://doi.org/10.1016/j.anbehav.2010.11.013

Vander Wal, E., Festa-Bianchet, M., Réale, D., Coltman, D. W., & Pelletier, F. (2015). Sex-based differences in the adaptive value of social behavior contrasted against morphology and environment. Ecology, 96(3), 631–641. https://doi.org/10.1890/14-1320.1

Vander Wal, E., Gagné-Delorme, A., Festa-Bianchet, M., & Pelletier, F. (2016). Dyadic associations and individual sociality in bighorn ewes. Behavioral Ecology, 27(2), 560–566. https://doi.org/10.1093/beheco/arv193

Villaret, J. C., & Bon, R. (1995). Social and spatial segregation in Alpine ibex (Capra ibex) in Bargy, French Alps. Ethology, 101(4), 291–300. https://doi.org/10.1111/j.1439-0310.1995.tb00366.x

Villaret J. C., Bon, R. (1998). Sociality and relationships in Alpine ibex (Capra ibex). Revue d’écologie.

von Hardenberg, A., Bassano, B., Arranz, M. D. P. Z., & Bogliani, G. (2004). Horn growth but not asymmetry heralds the onset of senescence in male Alpine ibex (Capra ibex). Journal of Zoology, 263(4), 425–432. https://doi.org/10.1017/S0952836904005485

Wasserman, S., & Faust, K. (1994). Social network analysis: Methods and applications (Vol. 8). Cambridge university press. https://doi.org/10.1017/CBO9780511815478

Welch, M. J., Smith, T., Hosie, C., Wormell, D., Price, E., & Stanley, C. R. (2020). Social Experience of Captive Livingstone’s Fruit Bats (Pteropus livingstonii). Animals, 10(8), 1321. https://doi.org/10.3390/ani10081321

Wey, T., Blumstein, D. T., Shen, W., & Jordán, F. (2008). Social network analysis of animal behaviour: a promising tool for the study of sociality. Animal behaviour, 75(2), 333–344. https://doi.org/10.1016/j.anbehav.2007.06.020

Wey, T. W., & Blumstein, D. T. (2010). Social cohesion in yellow-bellied marmots is established through age and kin structuring. Animal Behaviour, 79(6), 1343–1352. https://doi.org/10.1016/j.anbehav.2010.03.008

Wilson, A. D., Krause, S., James, R., Croft, D. P., Ramnarine, I. W., Borner, K. K., … & Krause, J. (2014). Dynamic social networks in guppies (Poecilia reticulata). Behavioral Ecology and Sociobiology, 68(6), 915–925. https://doi.org/10.1007/s00265-014-1704-0

Wilson, D. E., & Mittermeier, R. A. (2011). Handbook of the mammals of the word. Vol. 2: Hoofed mammals. Lynx Editions.

Wasserman, S., & Faust, K. (1994). Social network analysis: Methods and applications. Cambridge University Press, Cambridge (UK). https://doi.org/10.1017/CBO9780511815478

Whitehead, H. (2008). Analysing animal societies: Quantitative Methods for Vertebrate Social Analysis. Chicago: University of Chicago Press. https://doi.org/10.7208/chicago/9780226895246.001.0001

Whitehead, H. (2009). SOCPROG programs: analysing animal social structures. Behavioral Ecology and Sociobiology, 63(5), 765–778. https://doi.org/10.1007/s00265-008-0697-y

Willisch, C. S., & Neuhaus, P. (2009). Alternative mating tactics and their impact on survival in adult male Alpine ibex (Capra ibex ibex). Journal of Mammalogy, 90(6), 1421–1430. https://doi.org/10.1644/08-MAMM-A-316R1.1

Willisch, C. S., & Neuhaus, P. (2010). Social dominance and conflict reduction in rutting male Alpine ibex, Capra ibex. Behavioral Ecology, 21(2), 372–380. https://doi.org/10.1093/beheco/arp200

Willisch, C. S., Biebach, I., Koller, U., Bucher, T., Marreros, N., Ryser-Degiorgis, M. P., … & Neuhaus, P. (2012). Male reproductive pattern in a polygynous ungulate with a slow life-history: the role of age, social status and alternative mating tactics. Evolutionary Ecology, 26(1), 187–206. https://doi.org/10.1007/s10682-011-9486-6

Wittemyer, G., Douglas-Hamilton, I., & Getz, W. M. (2005). The socioecology of elephants: analysis of the processes creating multitiered social structures. Animal behaviour, 69(6), 1357–1371. https://doi.org/10.1016/j.anbehav.2004.08.018

Wittemyer, G., Okello, J. B., Rasmussen, H. B., Arctander, P., Nyakaana, S., Douglas-Hamilton, I., & Siegismund, H. R. (2009). Where sociality and relatedness diverge: the genetic basis for hierarchical social organization in African elephants. Proceedings of the Royal Society B: Biological Sciences, 276(1672), 3513–3521. https://doi.org/10.1098/rspb.2009.0941

