## Supplementary Material S0 for "Long term analysis of social structure: evidence of age-based consistent associations in male Alpine ibex"

**Table S0** Number of days of observation, number of surveys and of group observed, number of unique individuals and total number of animals observed during spring and summer season 2008-2017

| year | season | n days | n surveys | n groups observed | n ind unique | tot n ind observed |
| --- | --- | --- | --- | --- | --- | --- |
| 2008 | spring | 31 | 45 | 105 | 40 | 516 |
| 2008 | summer | 38 | 43 | 267 | 39 | 634 |
| 2009 | spring | 28 | 35 | 131 | 29 | 503 |
| 2009 | summer | 30 | 44 | 184 | 28 | 440 |
| 2010 | spring | 38 | 49 | 229 | 40 | 898 |
| 2010 | summer | 38 | 64 | 250 | 41 | 845 |
| 2011 | spring | 38 | 60 | 353 | 46 | 1293 |
| 2011 | summer | 36 | 57 | 508 | 44 | 1521 |
| 2012 | spring | 43 | 70 | 269 | 53 | 1659 |
| 2012 | summer | 27 | 39 | 363 | 49 | 1183 |
| 2013 | spring | 41 | 63 | 246 | 49 | 1428 |
| 2013 | summer | 41 | 58 | 266 | 46 | 821 |
| 2014 | spring | 35 | 53 | 280 | 40 | 1201 |
| 2014 | summer | 45 | 66 | 358 | 36 | 1201 |
| 2015 | spring | 44 | 68 | 255 | 47 | 1894 |
| 2015 | summer | 58 | 83 | 674 | 44 | 2015 |
| 2016 | spring | 31 | 43 | 148 | 39 | 1261 |
| 2016 | summer | 41 | 55 | 340 | 45 | 1463 |
| 2017 | spring | 39 | 68 | 344 | 42 | 1892 |
| 2017 | summer | 63 | 92 | 695 | 44 | 2300 |
