## Supplementary Material S1 for "Long term analysis of social structure: evidence of age-based consistent associations in male Alpine ibex"

The following pages contain the graphical representation of the weighted association networks of male Alpine ibex in spring and summer season of each year of the study (2008-2018).

Graphs were created using Gephi 0.9.2 (Bastian et al., 2009) with the "ForceAtlas" spatialization tool. The colours and width of the edges are proportional to the strength of association (with darker and thicker edges representing stronger associations). The size of the nodes represents the age of the individuals (due to space constrain on the page, the size of the nodes of animals of a given age is not constant in the graphs of different seasons/year). The colours of the nodes represent the value of the binary variable "season preceding death" of the individuals during the year. Individuals that die in the months following data collection are represented in darker colour.

### Alpine ibex network spring 2008

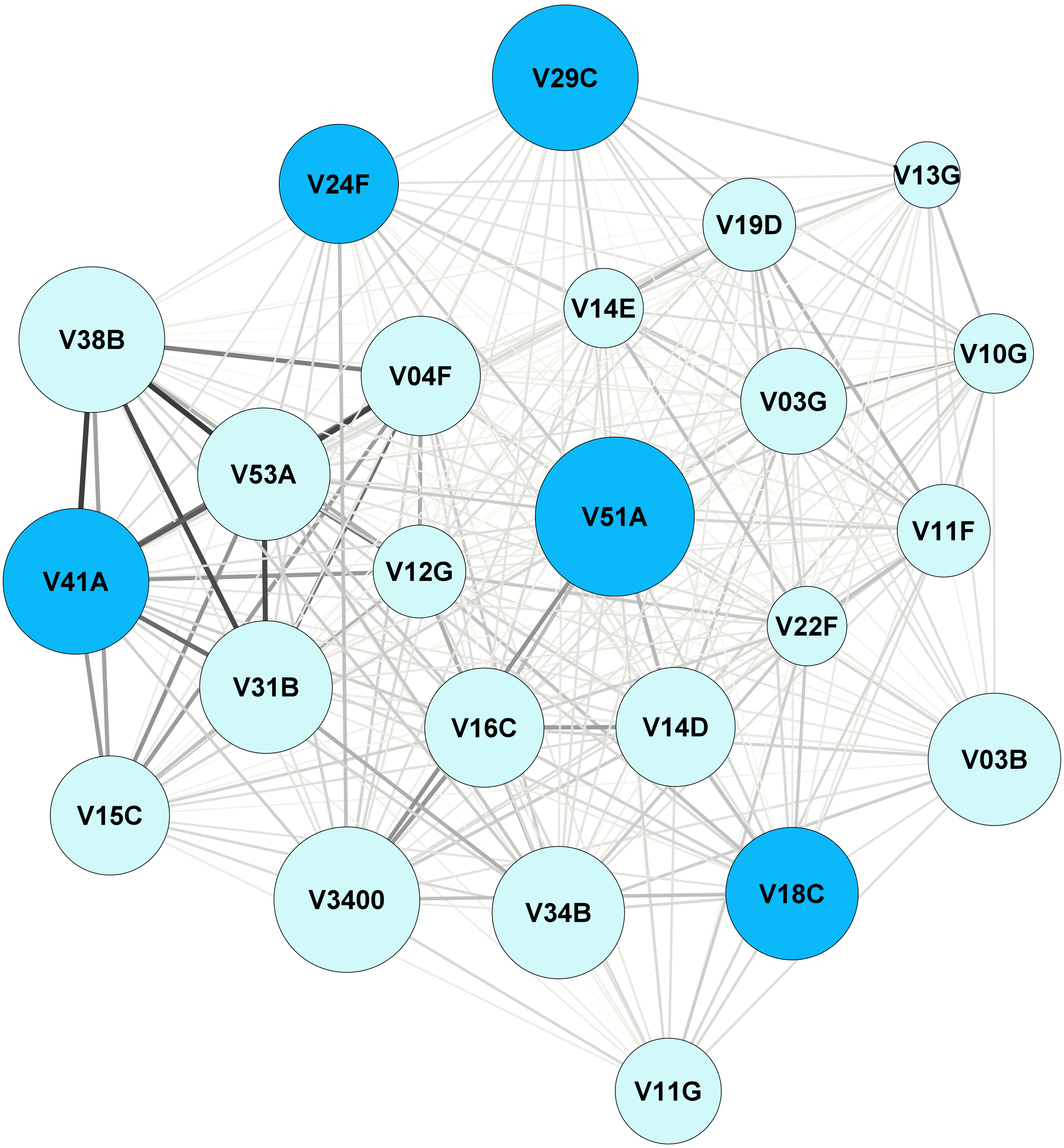

### Alpine ibex network summer 2008

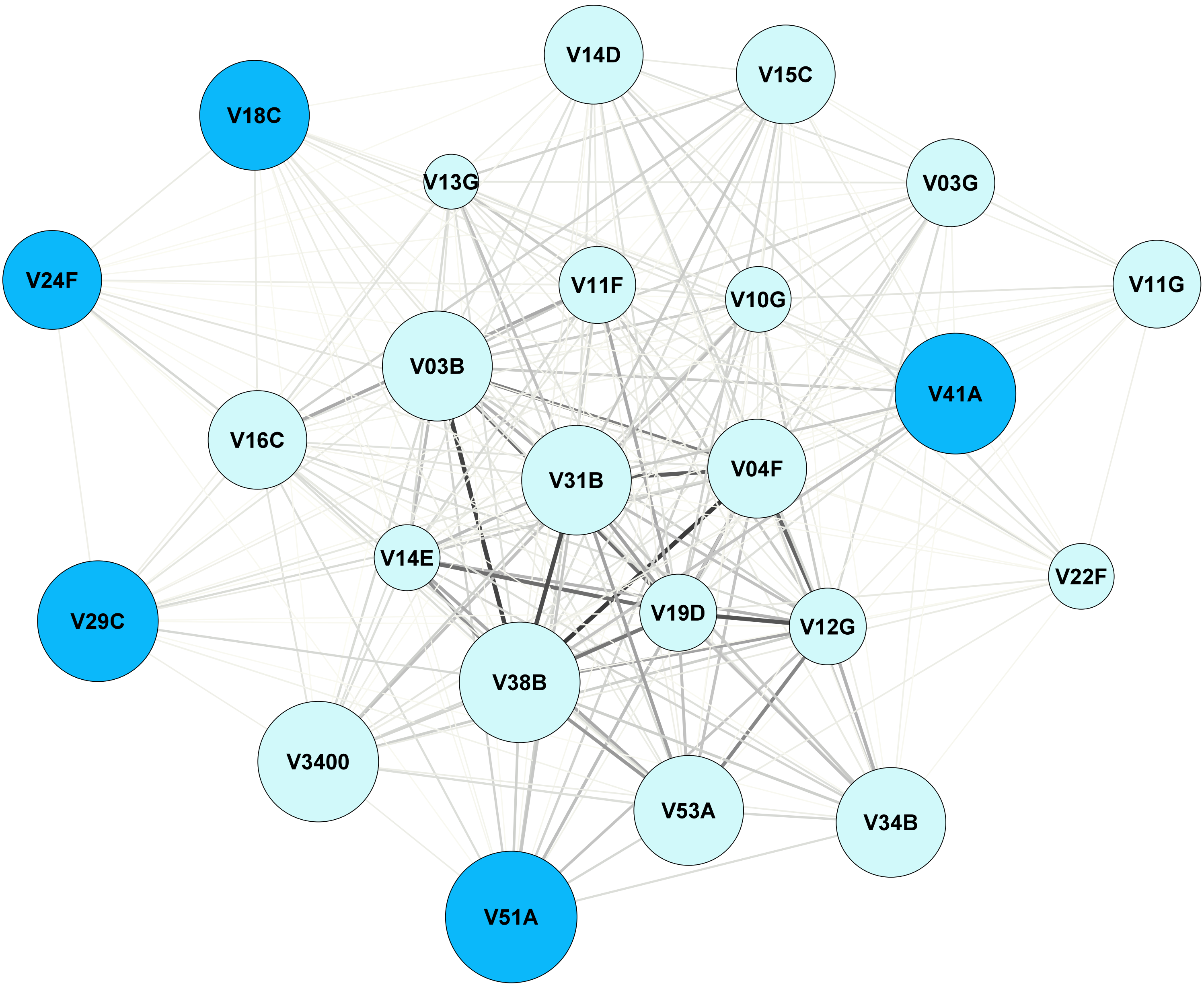

Alpine ibex network spring 2009

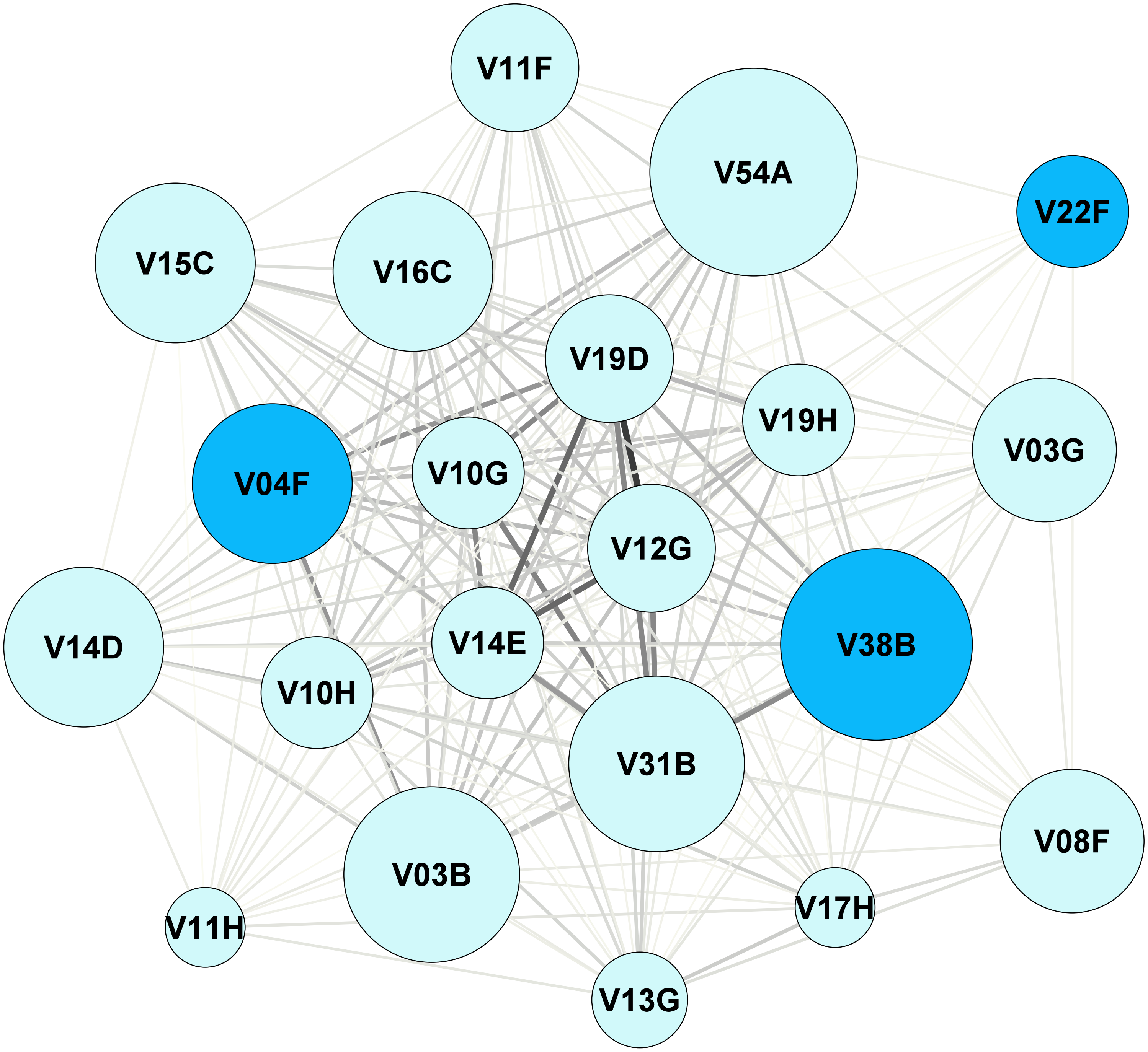

### Alpine ibex network summer 2009

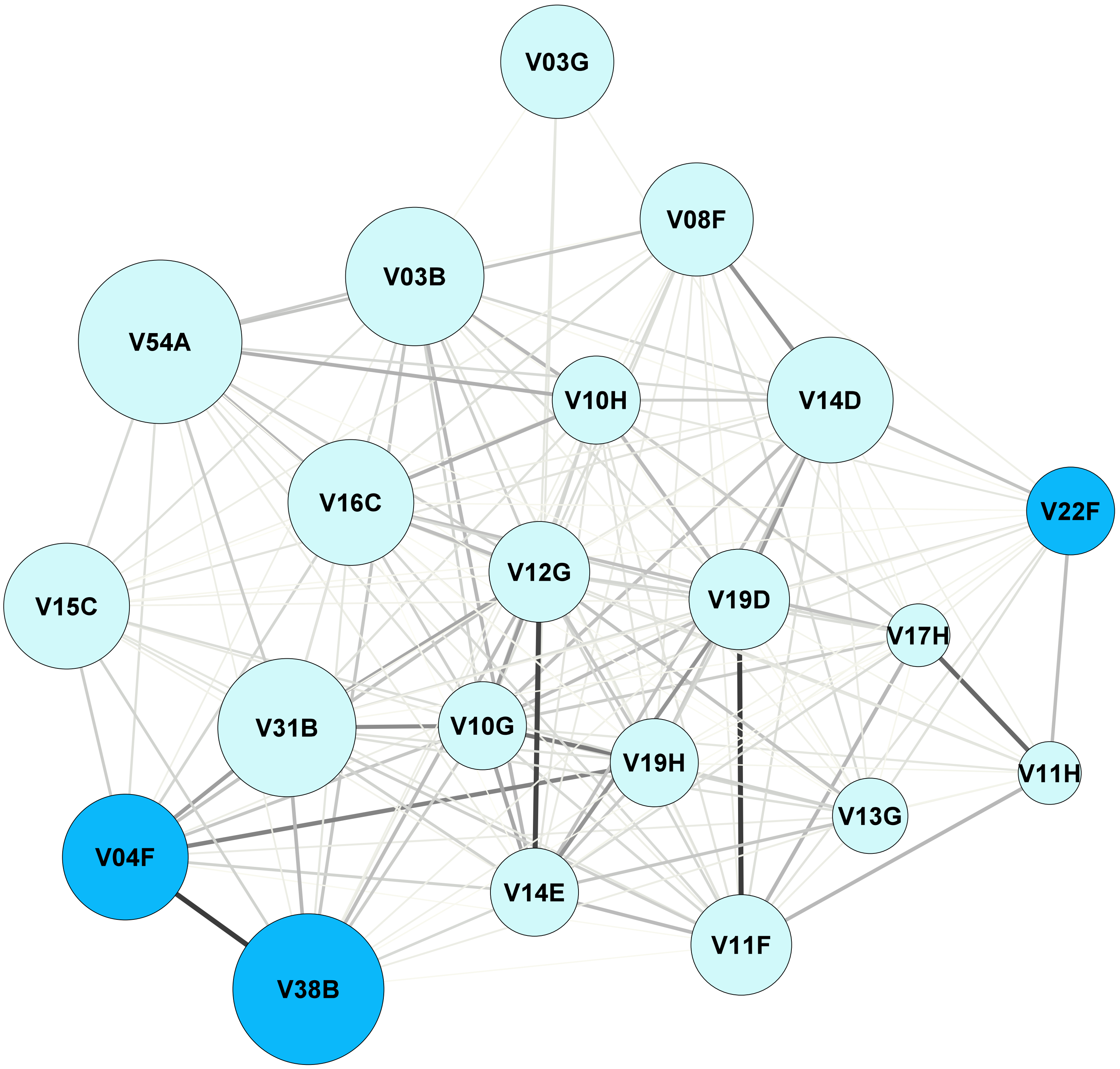

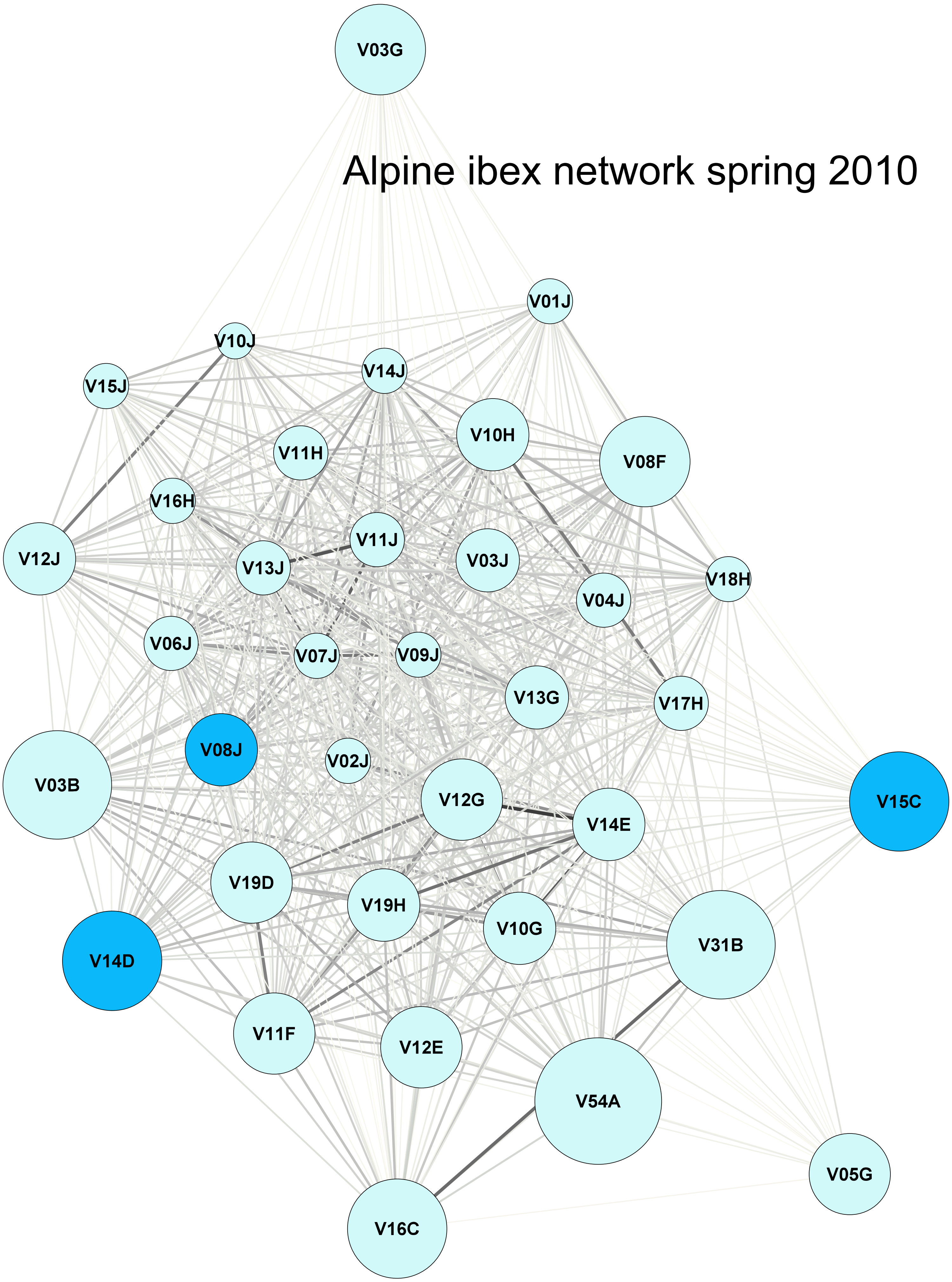
Alpine ibex network spring 2010
V03G
V01J
V10J
V14J
V15J
V10H
V11H
V08F
V16H
V11J
V12J
V13J
V03J
V18H
V04J
V06J
V07J
V09J
V13G
V17H
V08J
V02J
V03B
V15C
V12G
V14E
V19D
V19H
V10G
V31B
V14D
V11F
V12E
V54A
V05G
V16C

Alpine ibex network summer 2010

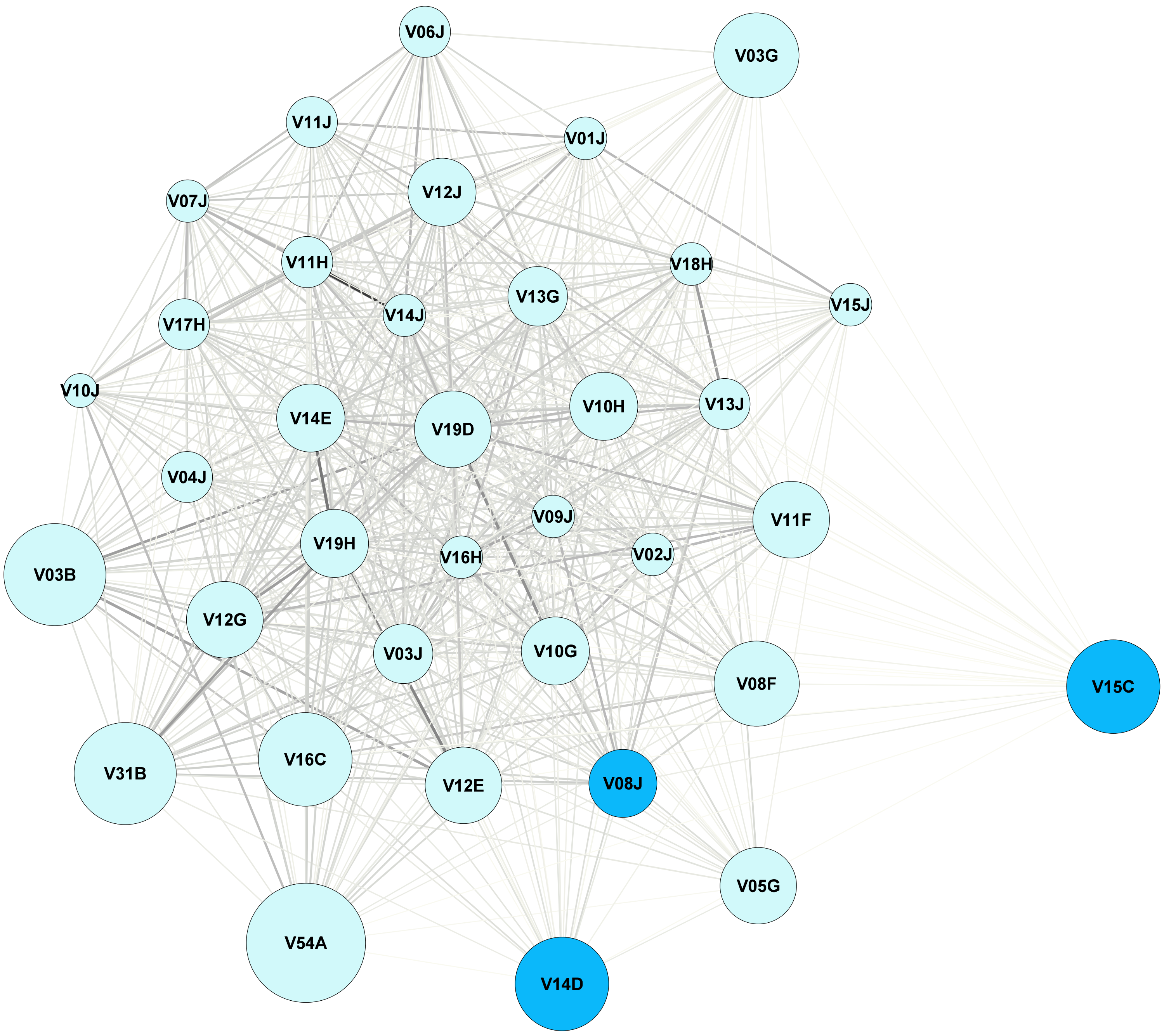

### Alpine ibex network spring 2011

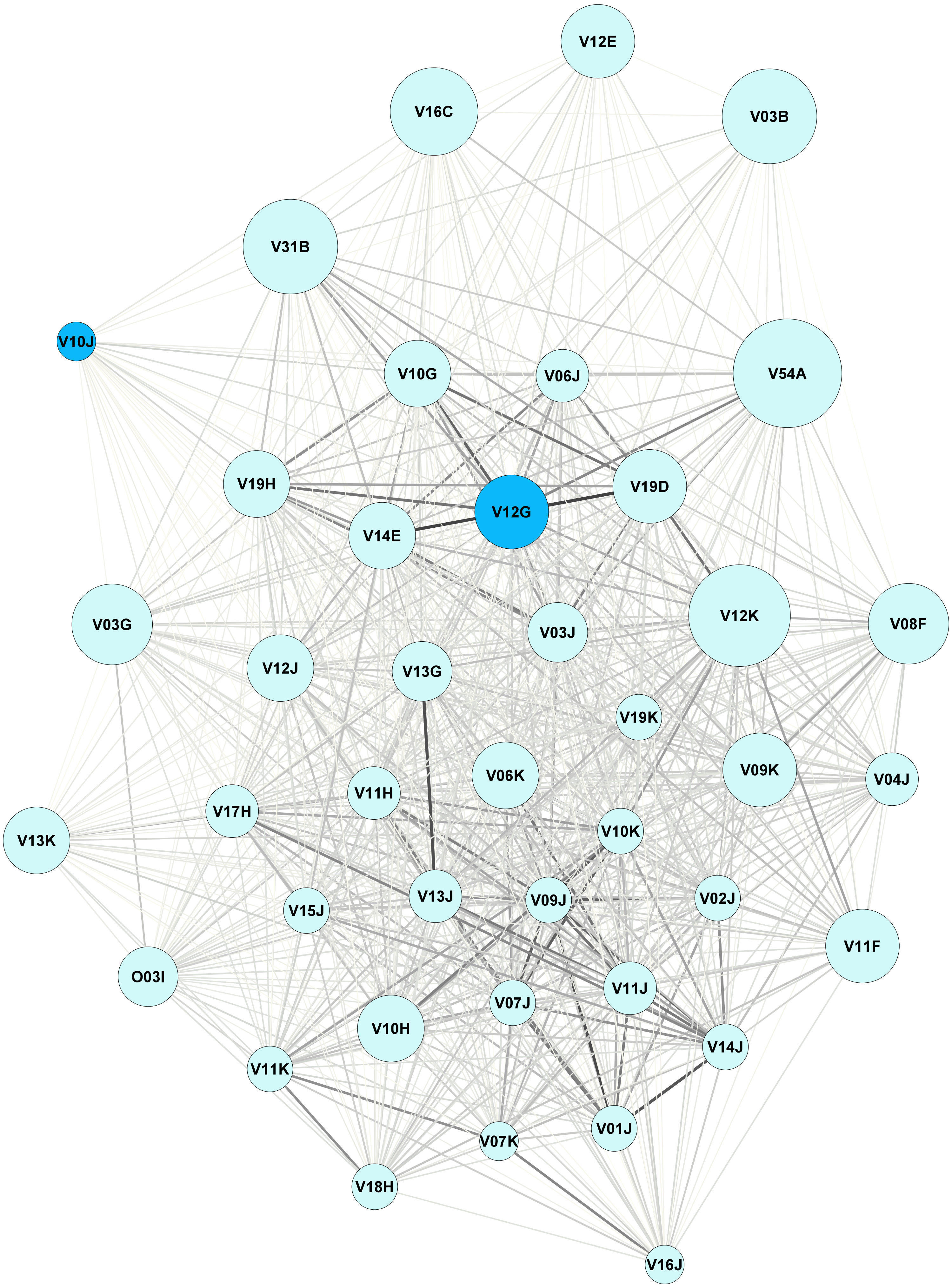

### Alpine ibex network summer 2011

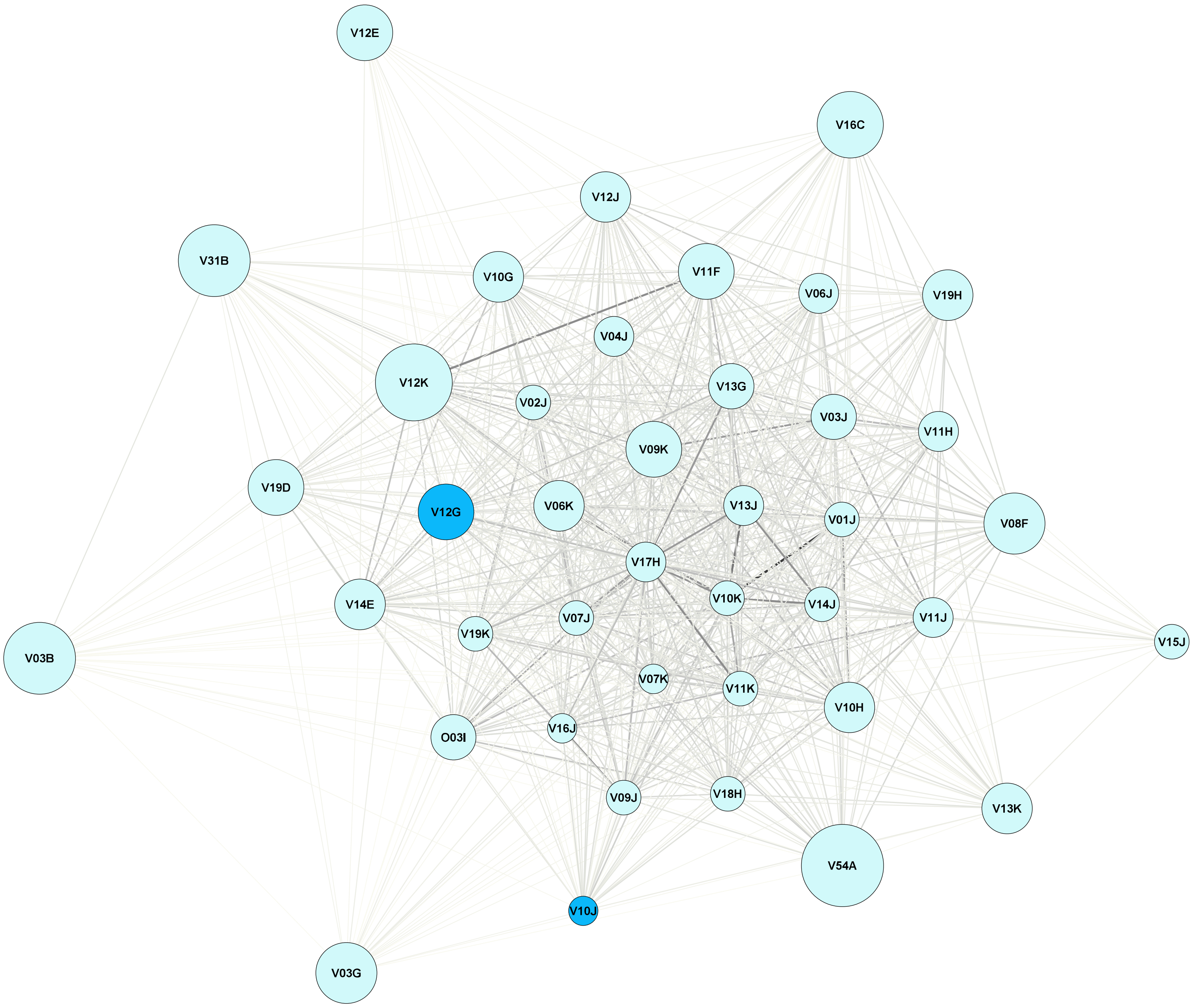

### Alpine ibex network spring 2012

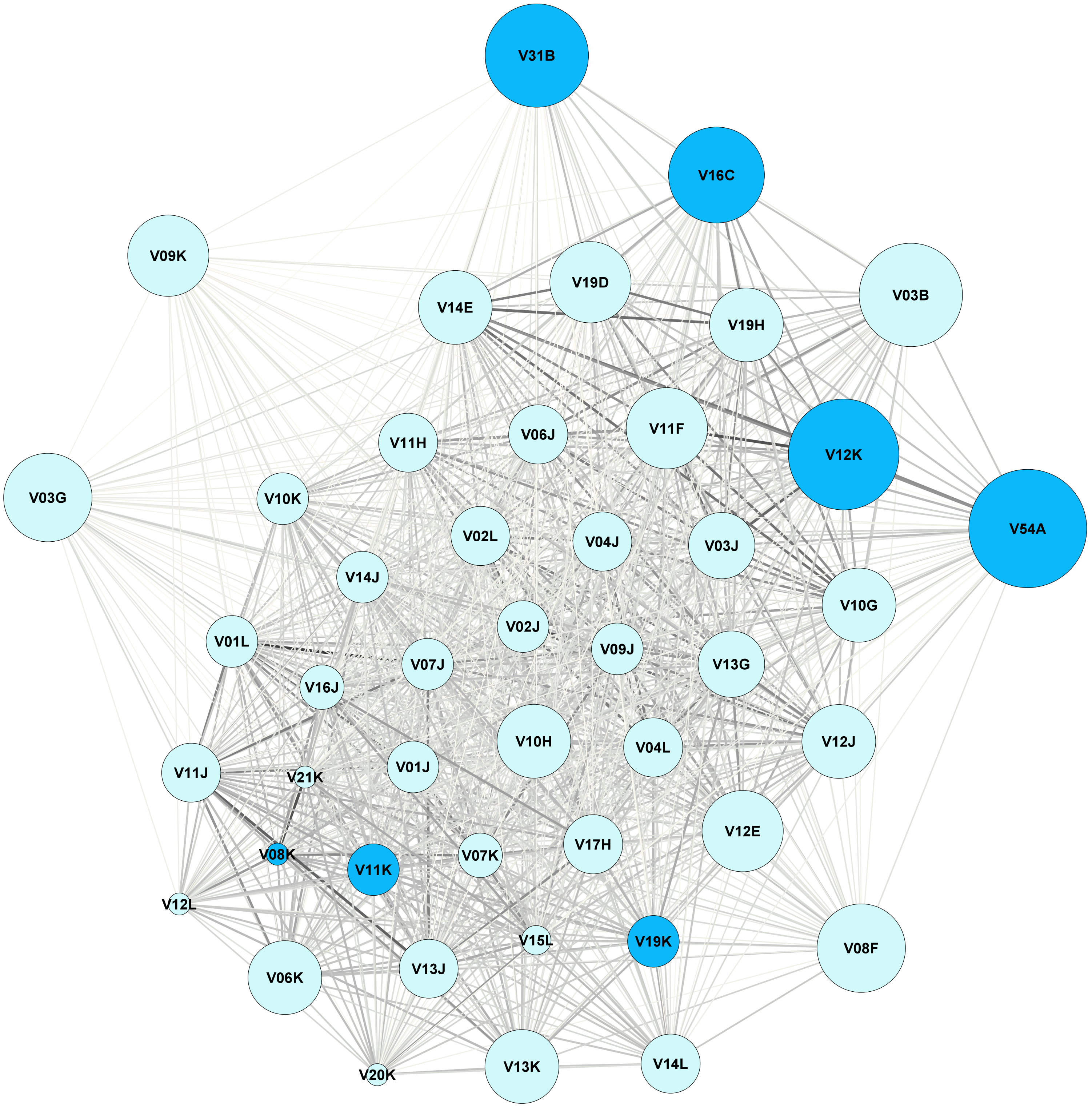

Alpine ibex network summer 2012

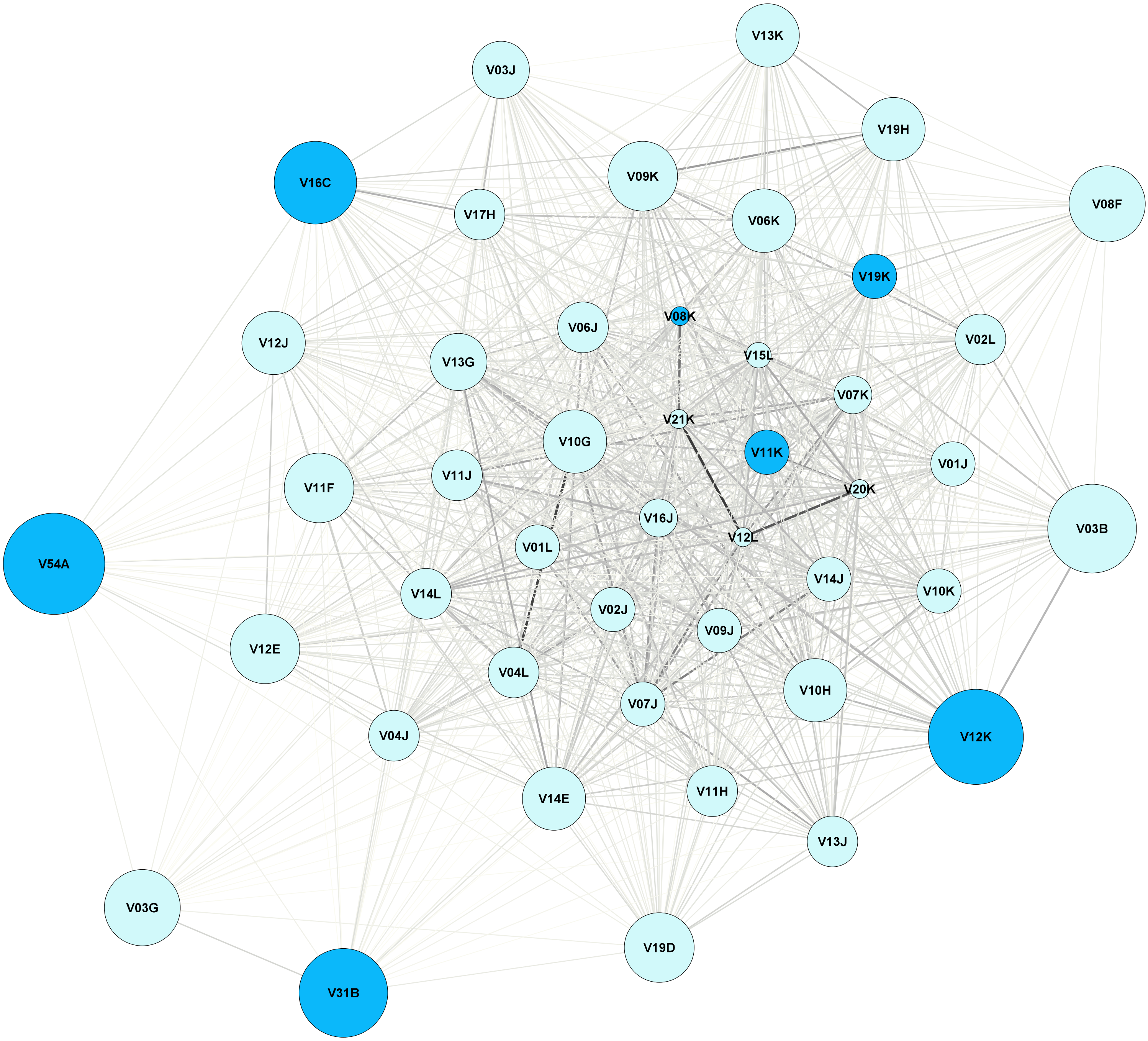

Alpine ibex network spring 2013

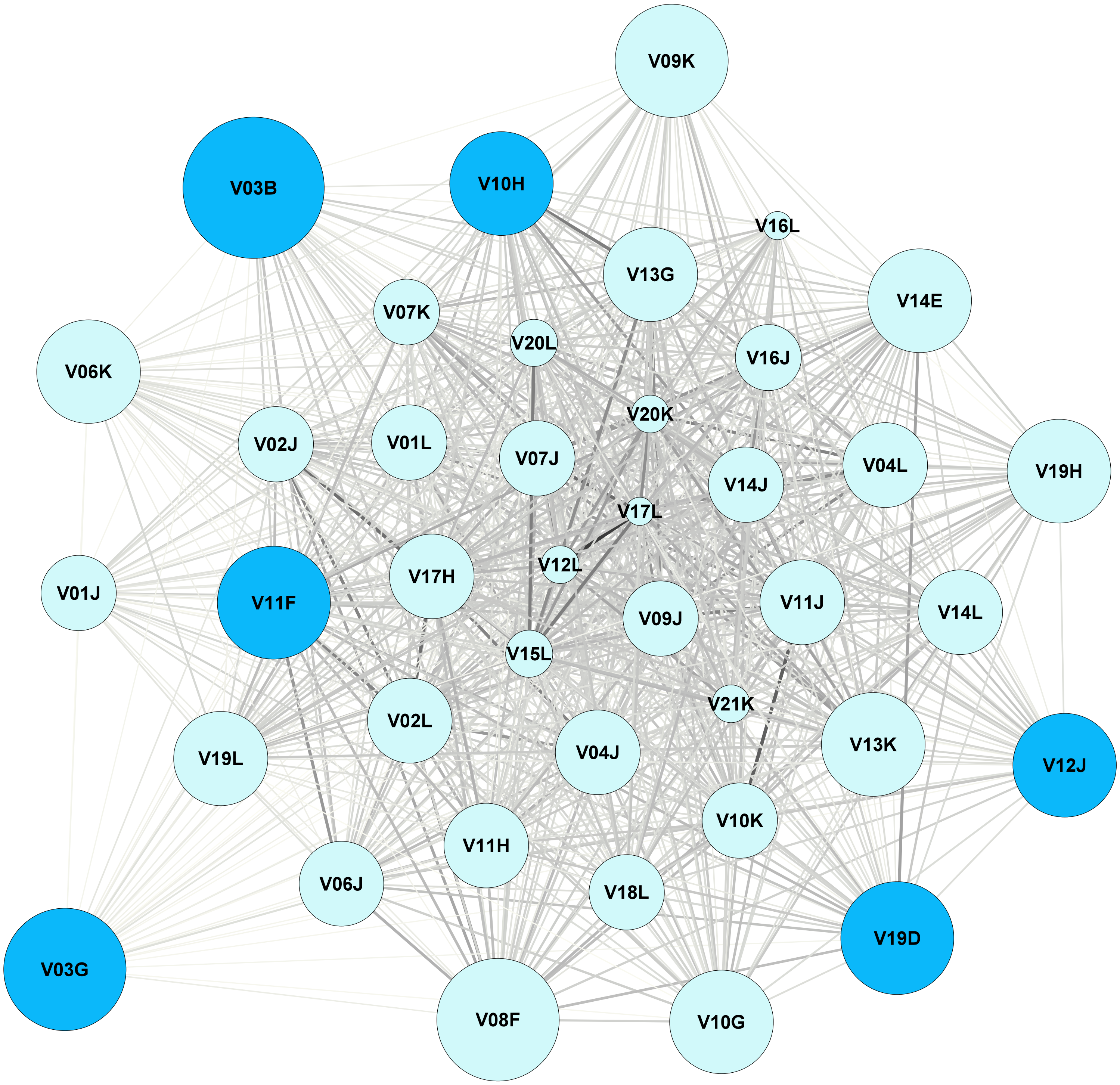

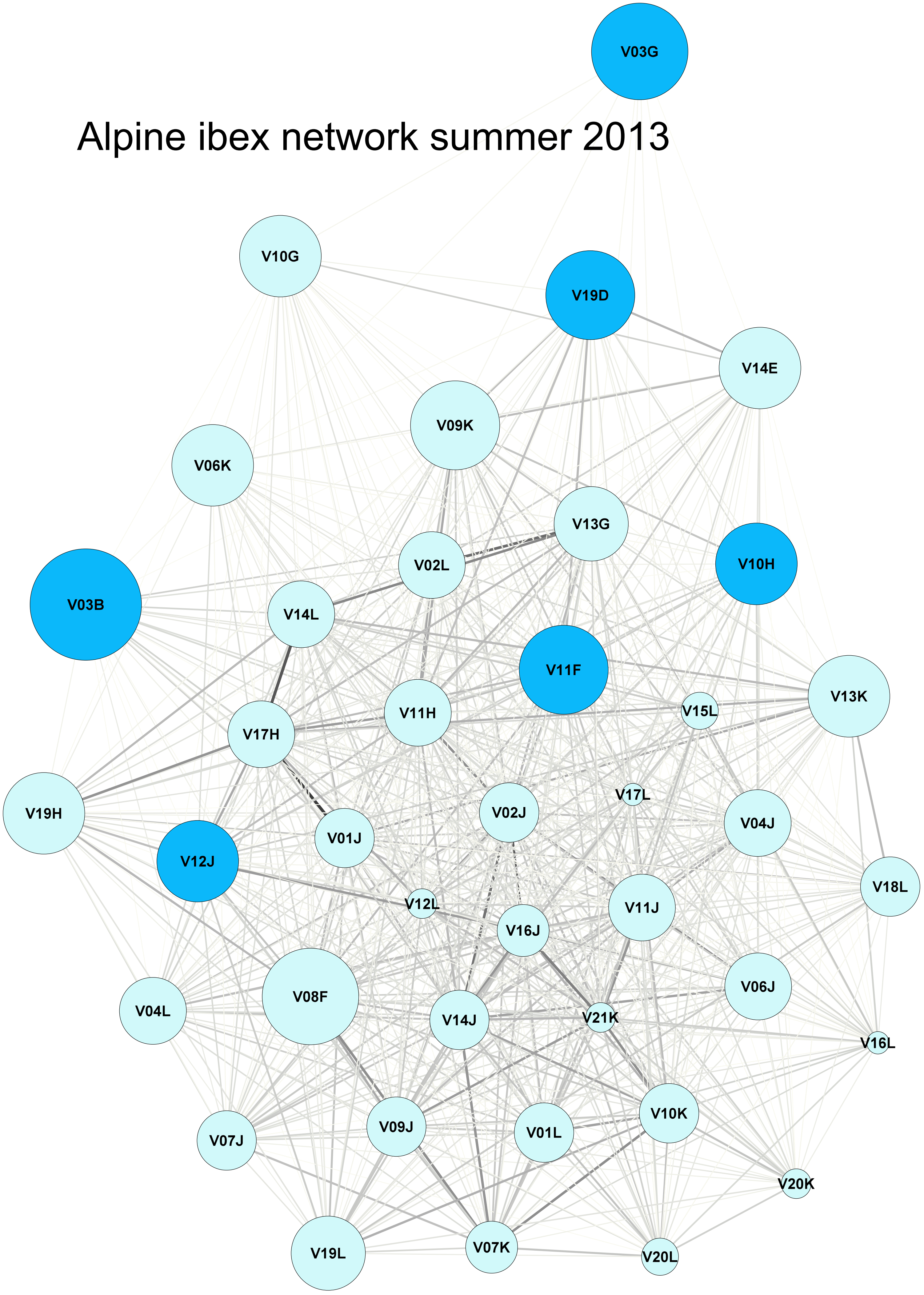
Alpine ibex network summer 2013
V03G
V10G
V19D
V14E
V09K
V06K
V13G
V02L
V10H
V03B
V14L
V11F
V13K
V11H
V15L
V17H
V17L
V19H
V02J
V04J
V01J
V12J
V18L
V12L
V11J
V16J
V06J
V08F
V04L
V14J
V21K
V16L
V10K
V09J
V01L
V07J
V20K
V07K
V19L
V20L

Alpine ibex network spring 2014

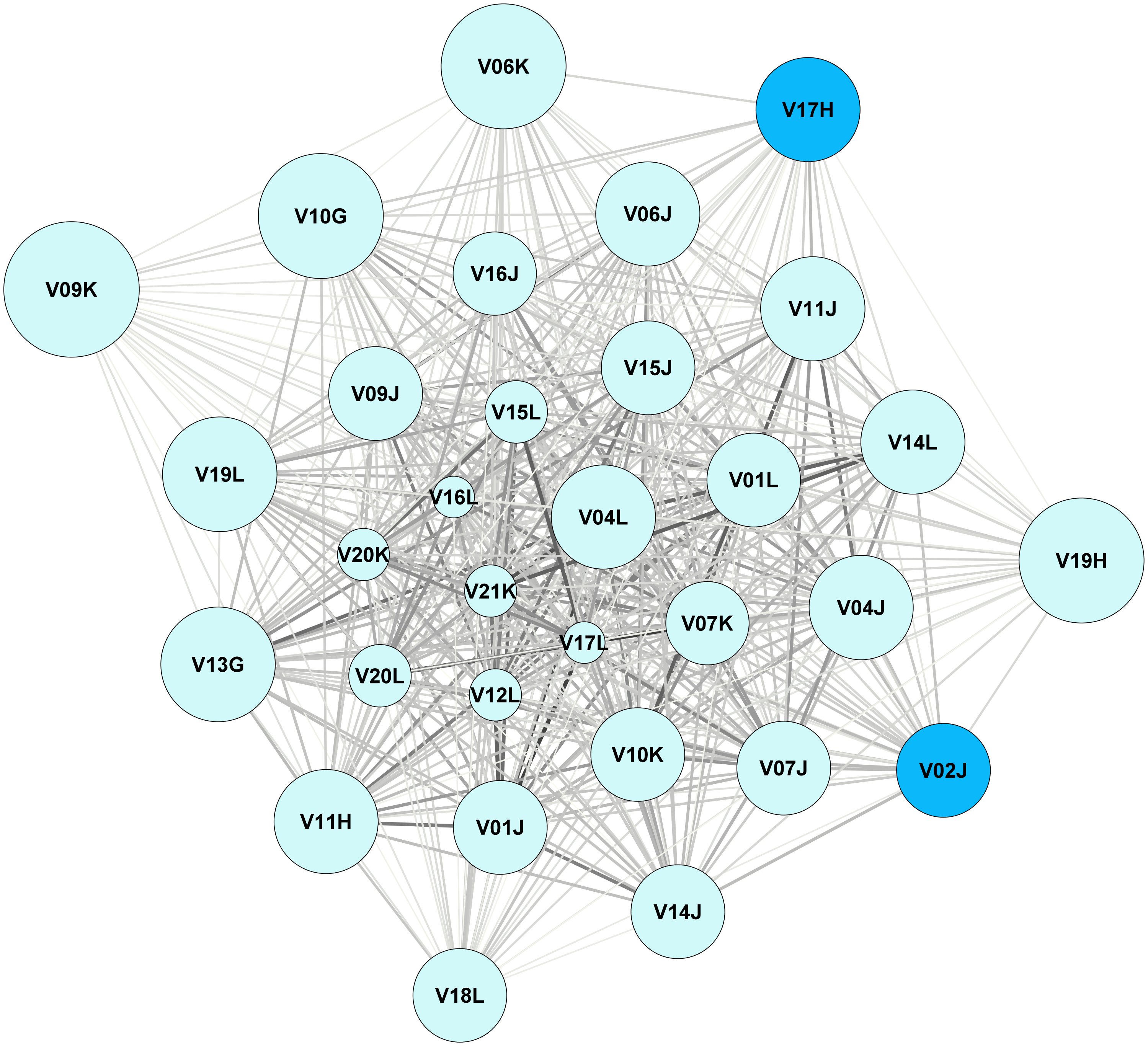

Alpine ibex network summer 2014

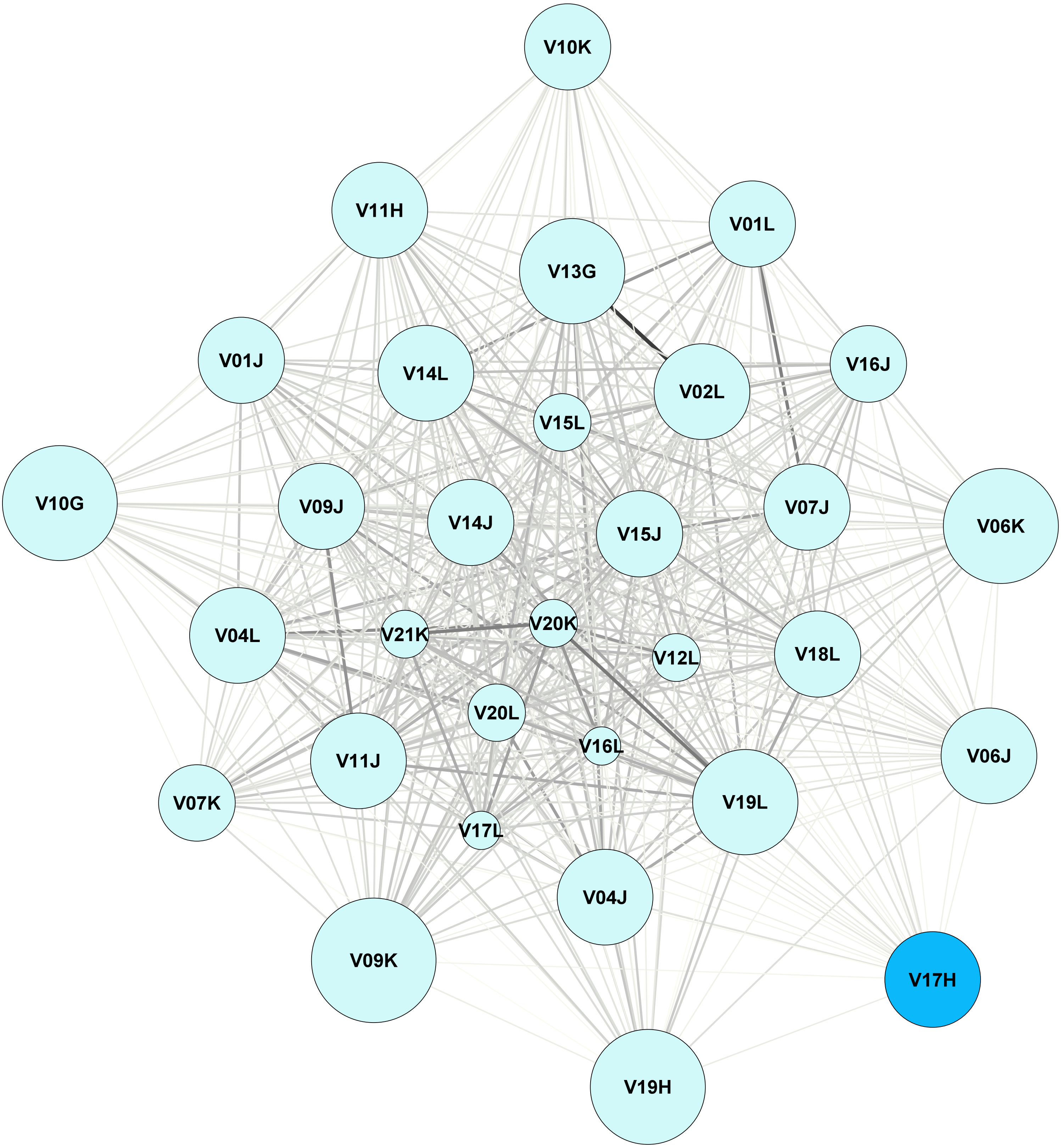

### Alpine ibex network spring 2015

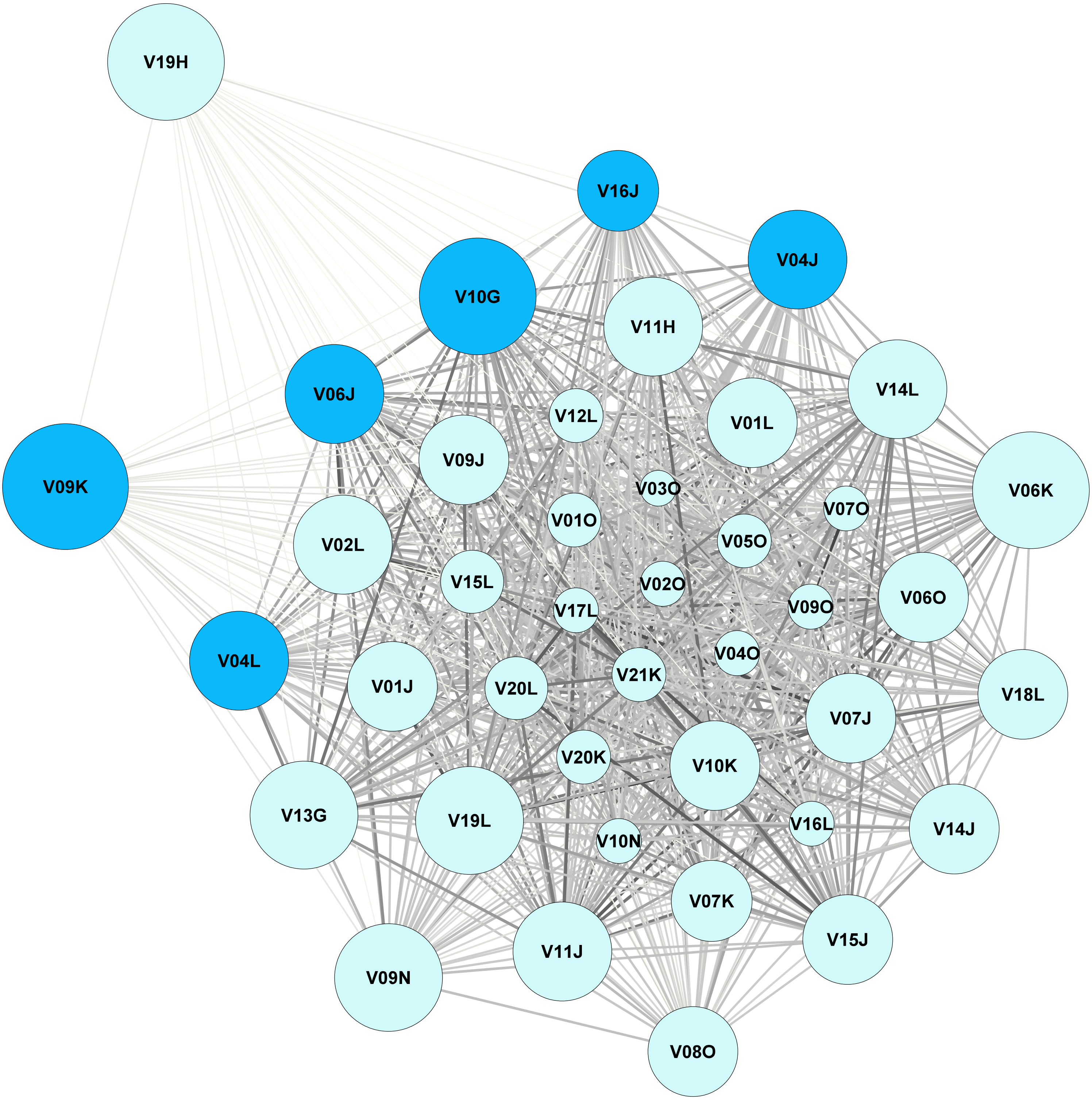

### Alpine ibex network summer 2015

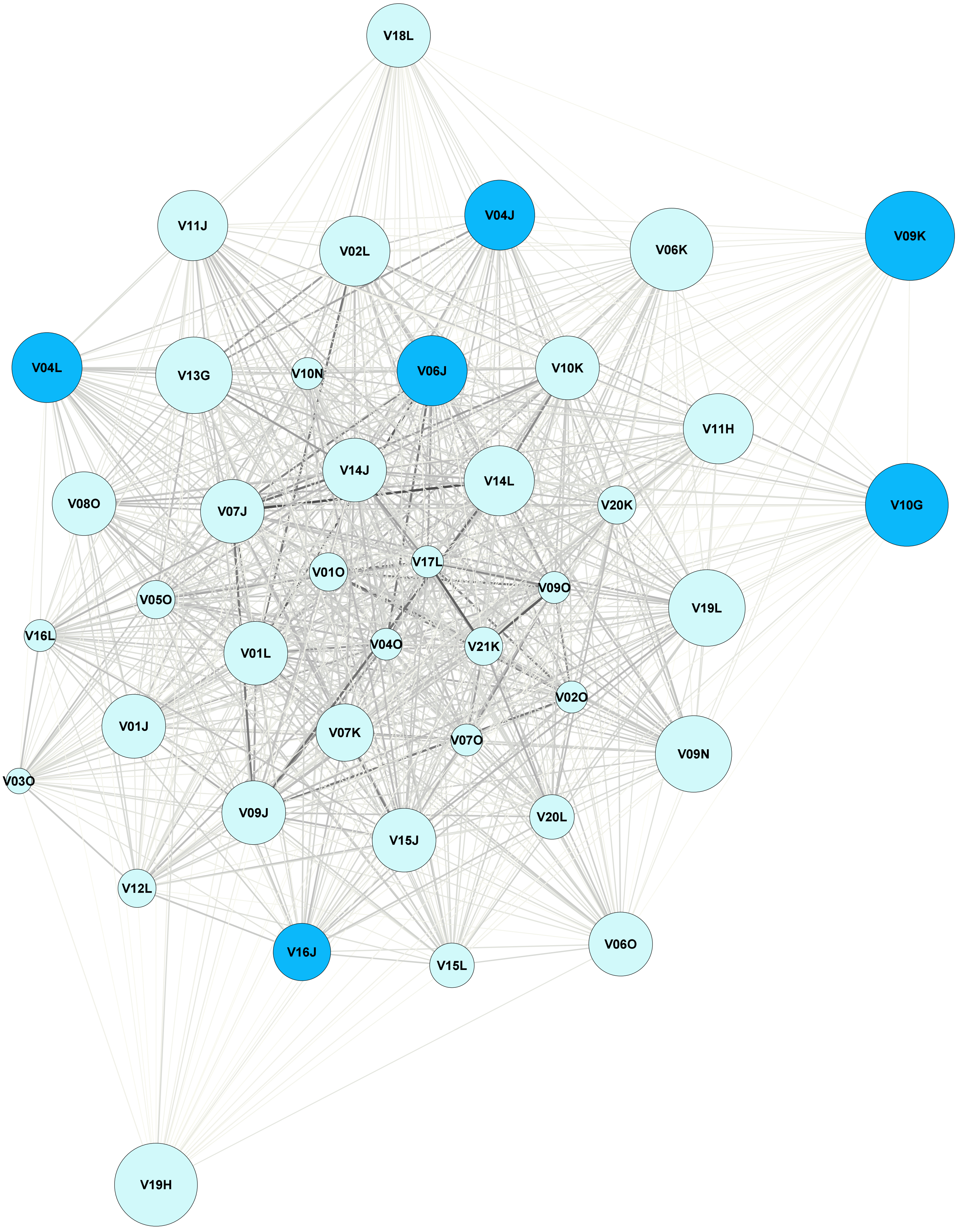

### Alpine ibex network spring 2016

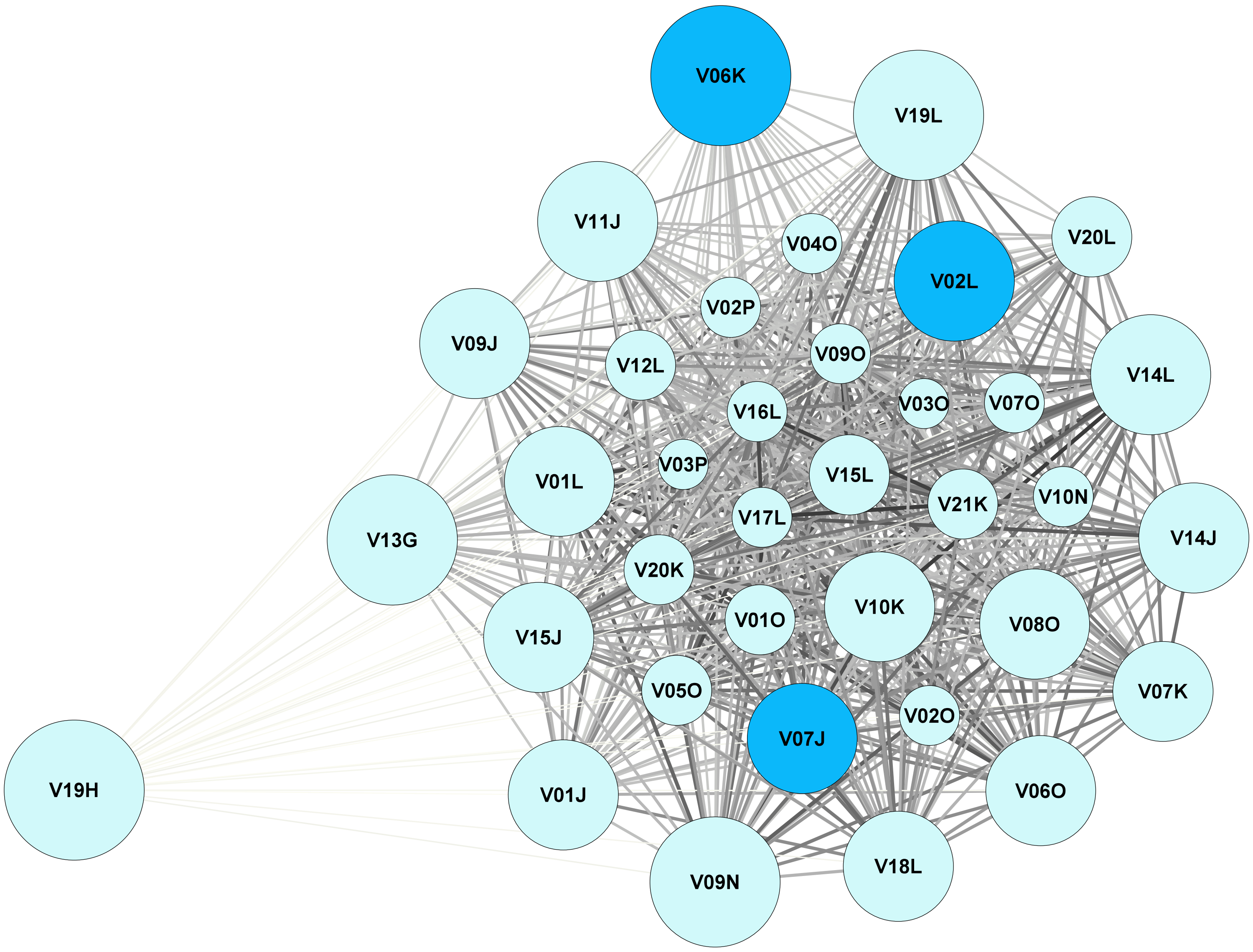

### Alpine ibex network summer 2016

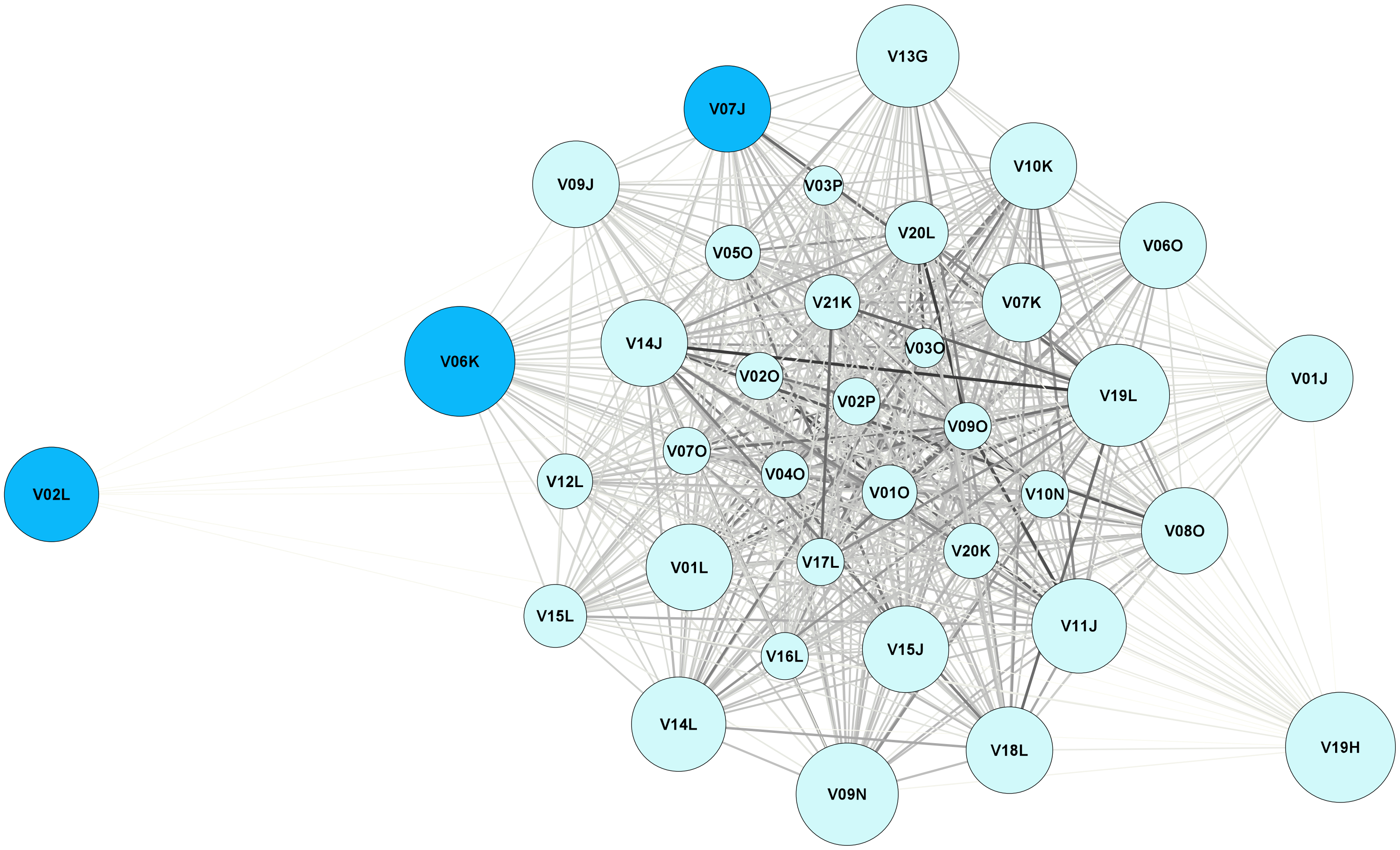

### Alpine ibex network spring 2017

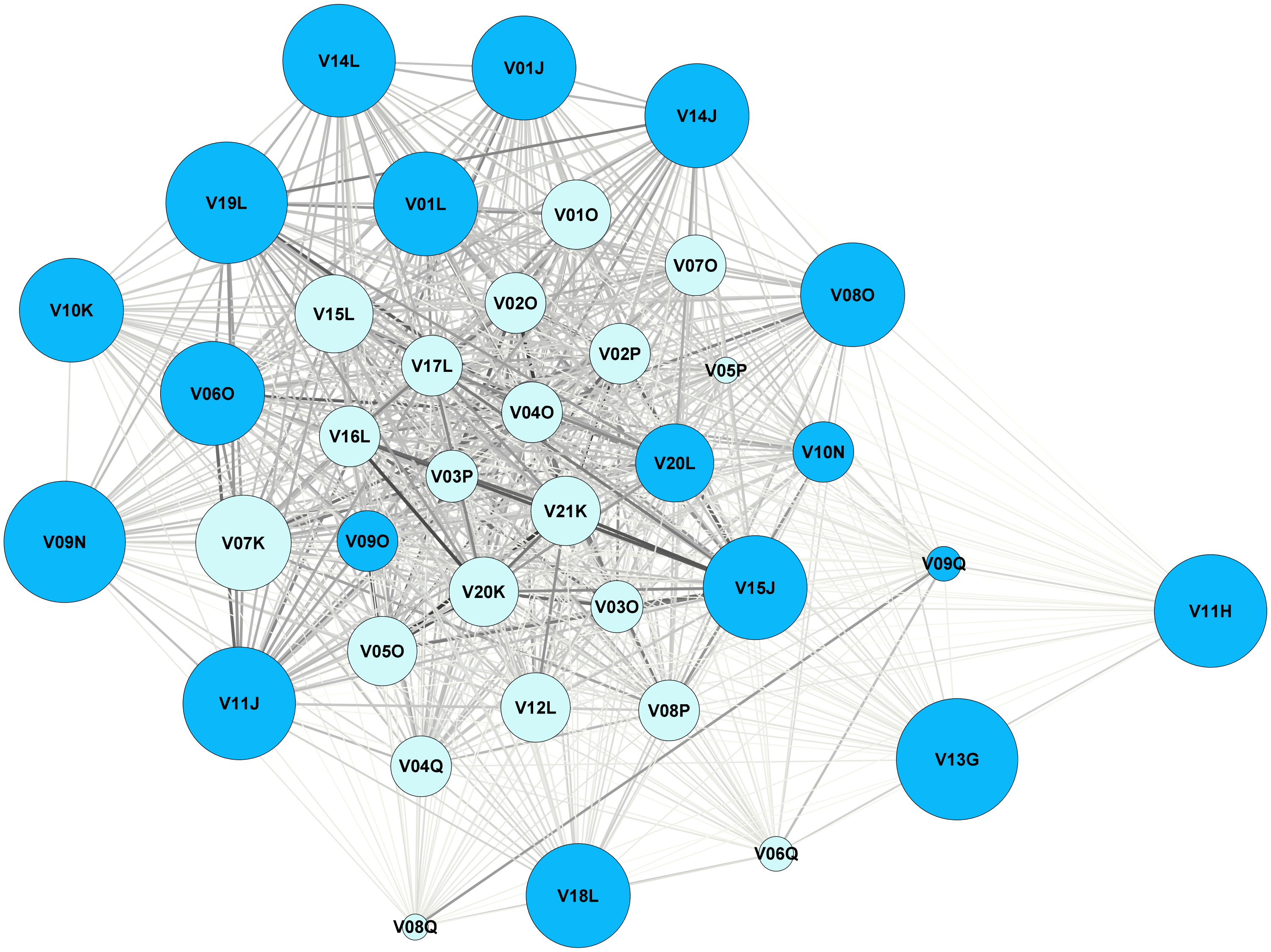

### Alpine ibex network summer 2017

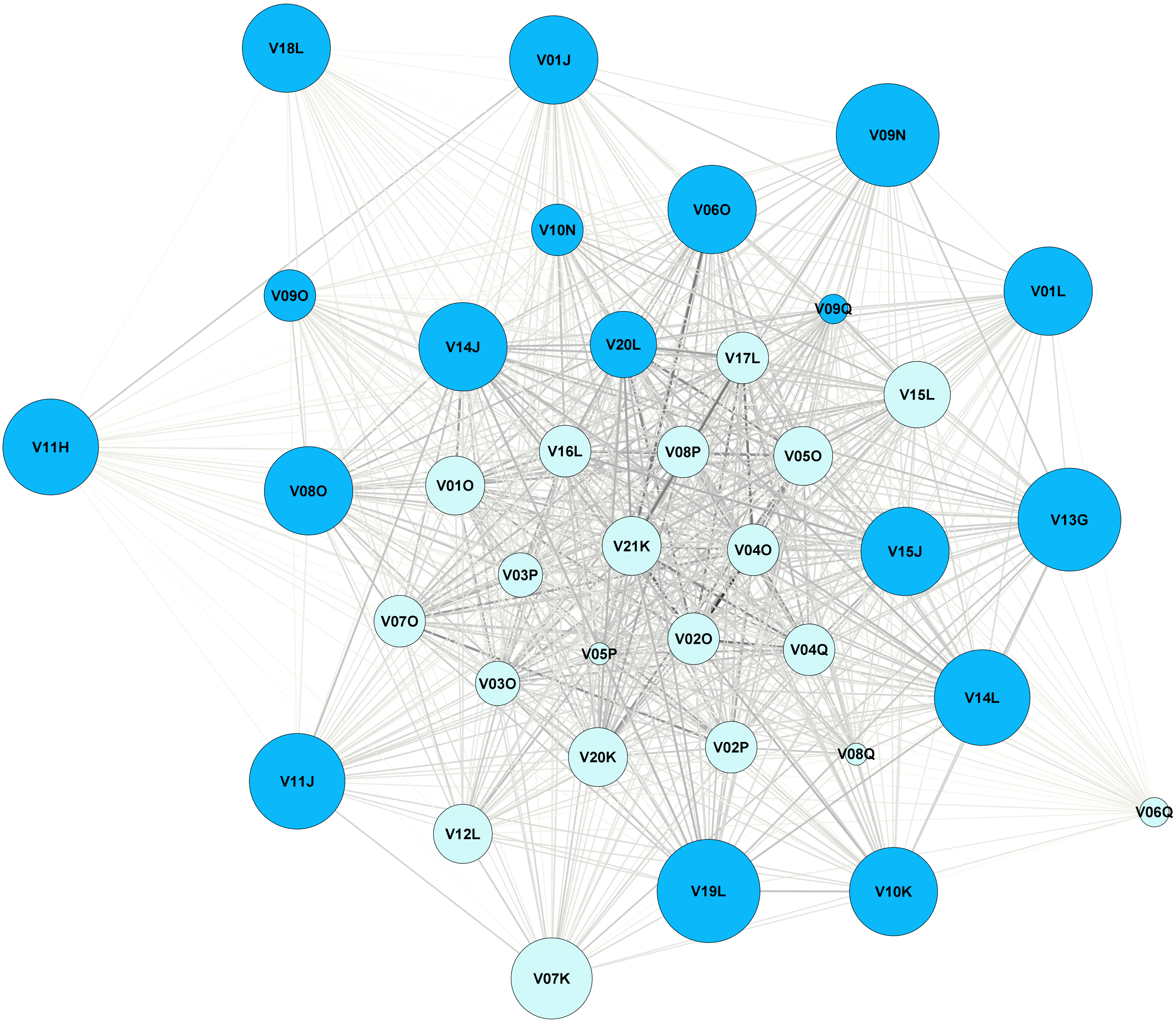
