## Supplementary Material S2 for "Long term analysis of social structure: evidence of age-based consistent associations in male Alpine ibex"

**Table S2a** Model selection for the GLMM performed to explain the variance of Strength Centrality

The table provides the fixed and random term included in the models, the degrees of freedom, the AIC and the ΔAIC compared to the best fitting model. The best fitting model is indicated in bold.

| N model | Fixed and random terms | *df* | AIC | ΔAIC to bfm |
| --- | --- | --- | --- | --- |
| 1 | age; age^2^; season; random: 1\|year; 1\|ID | 7 | 5330.80 | 16.25 |
| 1a | age; age^2^; season; random: 1\|year | 6 | 5457.42 | 142.87 |
| **2** | **age; age^2^; season prec. death; season; random: 1\|year; 1\|ID** | **8** | **5314.55** | **0** |
| 2a | age; age^2^; season prec. death; season; random: 1\|year | 7 | 5448.66 | 134.11 |
| 3 | age; age^2^; random: 1\|year; 1\|ID | 6 | 5815.38 | 500.83 |
| 3a | age; age^2^; random: 1\|year | 5 | 5840.18 | 525.63 |
| 4 | season; random: 1\|year; 1\|ID | 5 | 5408.27 | 93.72 |
| 4a | season; random= 1\|year | 4 | 5554.00 | 239.45 |

**Table S2b** Model selection for the GLMM binomial performed to explain the variance of Eigenvector Centrality. The table provides the fixed and random term included in the models, the degrees of freedom, the AIC and the ΔAIC compared to the best fitting model. The best fitting model is indicated in bold.

| N model | Fixed and random terms | *df* | AIC | ΔAIC to bfm |
| --- | --- | --- | --- | --- |
| 1 | age; age^2^; season; random: 1\|year; 1\|ID | 6 | 849.18 | 8.73 |
| 1a | age; age^2^; season; random: 1\|year | 5 | 860.89 | 20.44 |
| **2** | **age; age^2^; season prec. death; season; random: 1\|year; 1\|ID** | **7** | **840.45** | 0 |
| 2a | age; age^2^; season prec. death; season; random: 1\|year | 6 | 853.50 | 13.05 |
| 3 | age; age^2^; random: 1\|year; 1\|ID | 5 | 855.81 | 15.36 |
| 3a | age; age^2^; random: 1\|year | 5 | 866.01 | 25.56 |
| 4 | season; random: 1\|year; 1\|ID | 4 | 892.71 | 52.26 |
| 4a | season; random= 1\|year | 3 | 911.31 | 70.86 |
